# SwiftMHC: A High-Speed Attention Network for MHC-Bound Peptide Identification and 3D Modeling

**DOI:** 10.1101/2025.01.20.633893

**Authors:** Coos Baakman, Giulia Crocioni, Cunliang Geng, Daniel T. Rademaker, David Frühbuß, Yannick J. M. Aarts, Li C. Xue

## Abstract

Identifying tumor peptides that bind patient MHC proteins and elicit immune responses is central to immunotherapy, yet progress remains limited to a handful of well-studied alleles. Structure-based methods generalize better than sequence-only models but are constrained by the high computational cost of 3D modeling. We present SwiftMHC, the fastest structure-based framework for peptide–MHC (pMHC) modeling and binding affinity prediction. SwiftMHC predicts peptide–MHC binding in 0.009 sec/case in batch mode on a single A100 GPU—nearly an order of magnitude faster than leading sequence-based methods such as NetMHCpan 4.1 (0.081 sec/case)—while performing competitively in predictive accuracy. In addition to affinity estimation, SwiftMHC generates *all-atom* 3D pMHC structures and achieves a median Cα-RMSD of 1.32 Å against X-ray benchmarks, matching or better than the accuracy of state-of-the-art approaches (e.g., AlphaFold with fine-tuning) but running thousands of times faster (excluding disk-writing time). These results demonstrate the power of task-specific AI trained on physics-derived synthetic data to overcome the scarcity of experimental structures. Optimized for HLA-A*02:01 9-mers but readily extensible to other alleles, SwiftMHC enables rapid and accurate identification of peptides distinct from self at T-cell–exposed surfaces. This capability may expand immunotherapy targets, improve the safety of TCR-based therapies, and accelerate the development of next-generation cancer immunotherapies.

## Main

Advances in understanding T cell immunity have led to breakthroughs in cancer immunotherapy, where the immune system is trained to recognize and destroy cancer cells, offering a promising alternative to traditional treatments with fewer side effects^1^. T cells are activated when their T-cell receptor (TCR) recognizes tumor-specific peptides presented on the tumor cell surface by the class I major histocompatibility complex (MHC-I) proteins (**Fig. 1A** for molecular details). However, these therapies face challenges, including high costs, long development time and toxicity risks^1,2^. Currently, the limited number of available target peptides is one of the major bottlenecks of cancer immunotherapies^3^.

**Figure 1.**
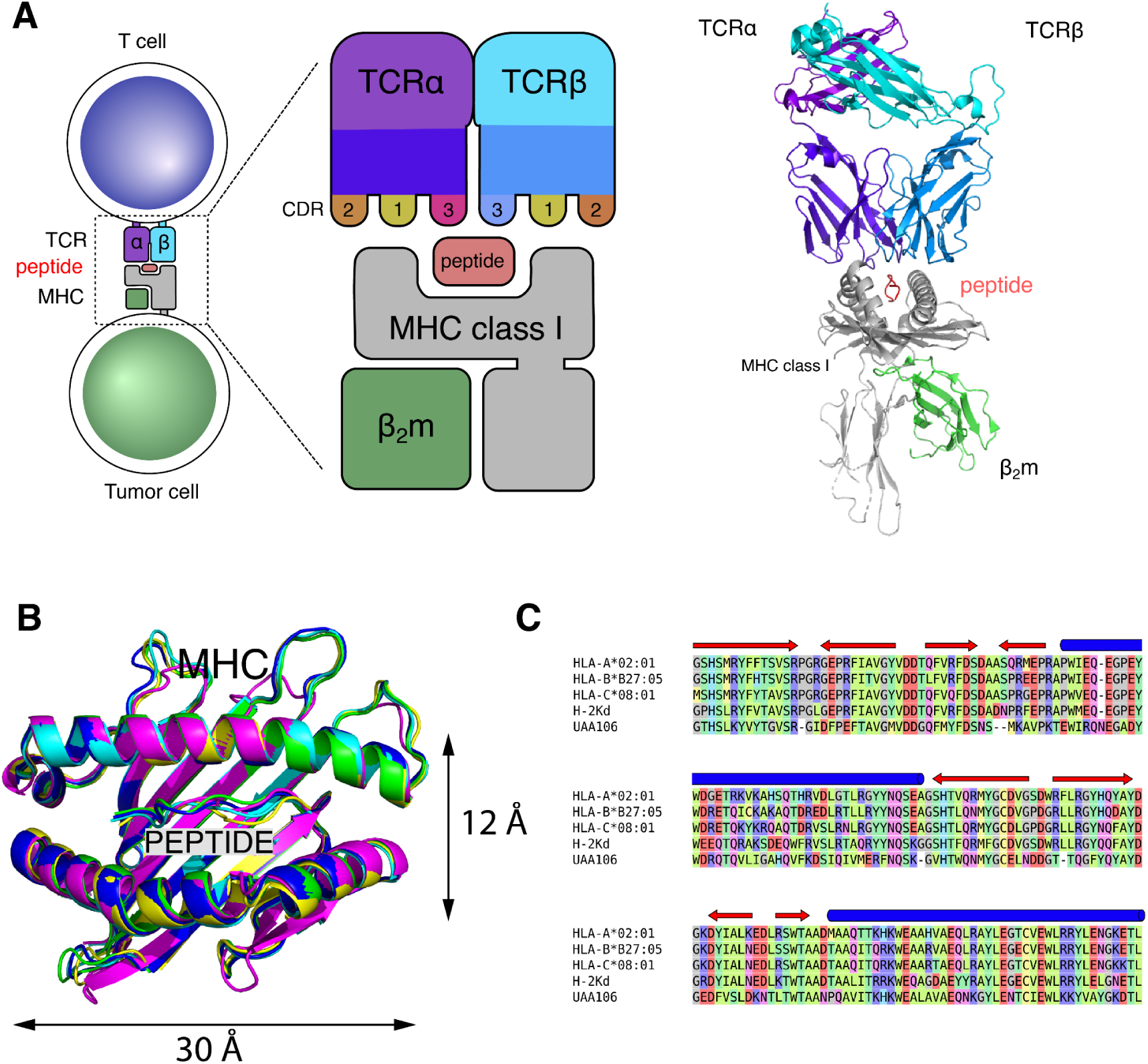
The TCR-peptide-MHC class I (TCR:pMHC-I) complex and neoantigens: their central role in immune surveillance and T-cell-mediated immune attacks on tumor cells. **A.** TCR nomenclature and the TCR:pMHC-I complex. A TCR has two chains (α and β chains), each having three loops (CDR1, CDR2, and CDR3), where CDR3 plays the primary role in interacting with the peptide. The image on the right shows the structural representation of a TCR:pMHC-I complex based on PDB entry 7RTR. **B.** Structural alignment of five different MHC alleles: HLA-A*02:01 (Homo sapiens, PDB ID: 5HHN), HLA-B*27:05 (Homo sapiens, PDB ID: 5IB2), HLA-C*08:01(Homo sapiens, PDB ID: 4NT6), H-2Kd (Mus musculus, PDB ID: 1VGK), and UAA106 (Ctenopharyngodon idella, bony fish, PDB ID: 6LBE). The MHC G-domains all consist of two ɑ-helices and a β-sheet. The peptides are bound in the MHC binding groove. The root mean square deviation (RMSD) values for the structural alignment of these G-domains are all below 0.8 Å, demonstrating highly conserved MHC structures across different alleles and species. **C.** Sequence alignment of MHC G-domains. The sequence alignment of the MHC G-domains from the five alleles shown in panel B demonstrates high diversity in the MHC sequences. The alignment was generated using Clustal Omega^20^ and visualized in MRS^21^ with colors representing the charge, hydrophobicity, size and shape of amino acid side chains. Blue cylinders above the sequences indicate α-helices, while red arrows denote β-strands.

Many predictive methods have been developed to identify MHC-binding peptides^4,5^, which have recently already made significant contributions in cancer immunotherapies design^6^. State-of-the-art (SOTA) tools, such as MHCflurry 2.0^7^ and netMHCpan 4.1^8^ use both the sequence of the MHC binding groove and the peptide sequence as input. However, these sequence-based approaches have notable limitations: they require large training datasets due to their reliance on sequence information and have limited generalizability on unseen alleles^9^; they overlook critical structural features of peptide-MHC interactions; and they struggle to handle peptides of varying lengths. Moreover, they do not provide 3D structural models, which are essential for understanding immunogenicity and guiding TCR design.

Alternatively, 3D structure-based approaches offer several compelling advantages: 1) they naturally handle peptide length variability in 3D space; 2) they are sensitive to mutations in both the spatial and energy landscapes; 3) they are potentially more robust for rare alleles due to the high conservation of MHC structures^9^ (**Fig. 1BC**); 4) 3D structures provide fundamental insights into the mechanisms of immunotherapies. However, experimental methods like X-ray crystallography, NMR and cryo-EM are labor-intensive and cannot keep up with the diversity of pMHCs. To date, only ∼1000 pMHC structures are available in the Protein Data Bank (PDB, www.rcsb.org)^10^, as contrast to the high diversity of human leukocyte antigens (HLAs, over 40,000 variancts identified)^11^. Therefore, complementary 3D modeling techniques are valuable tools.

Physics-based homology modeling tools such as PANDORA^12^ and APE-Gen 2.0^13^ can generate multiple MHC-I/II models, and provide energy scores indicative of binding affinity (BA). Yet, they remain computationally expensive (seconds to minutes per case). AlphaFold^14,15^, a powerful deep learning framework for 3D modeling, also faces limitations: it relies on computationally heavy modules to process multiple sequence alignments (MSAs), which are often unavailable for short peptides lacking evolutionary depth. AlphaFold2-FineTune^16^ a variant adapted for pMHC BA prediction, bypasses MSA searches by using query-to-template alignments, but still depends on the costly Evoformer module—, a limitation also reported by Mikhaylov et al. ^17^

Other methods attempt to address these challenges. MHCfold uses convolutional neural networks to predict 3D pMHC structures and then use multi-headed attention on the predicted structures to predict BAs. However, it requires additional tools (e.g., MODELLER^18^ or SCWRL4^19^) for side-chain reconstruction. Importantly, recent studies^9^ show that 3D model–based BA predictors have better generalizability than sequence-only methods on unseen alleles (6-16%), but they typically follow a two-step pipeline: (i) generating 3D pMHC models (seconds–minutes per case), and (ii) applying geometric deep learning (GDL) networks for BA estimation. This workflow is time-consuming and poorly scalable, particularly for large-scale screening of patient-derived mutations. Thus, there is a clear need for a time-efficient system capable of simultaneously predicting 3D structures and BA in a single step.

Here, we present SwiftMHC, a transformer that simultaneously predicts pMHC-I BA and generates corresponding 3D structures in milliseconds (**Fig. 2A**). By leveraging the conserved architecture of MHC proteins and avoiding computationally intensive MSA-based modules like AlphaFold’s Evoformer, SwiftMHC ensures rapid and accurate predictions. To address the challenge of modeling peptide conformations without evolutionary constraints, we enhance the training dataset with physics-based 3D models and biochemistry-derived BA data, enabling synergistically improvements in both structural and affinity predictions through residue-to-residue attention mechanisms.

**Figure 2.**
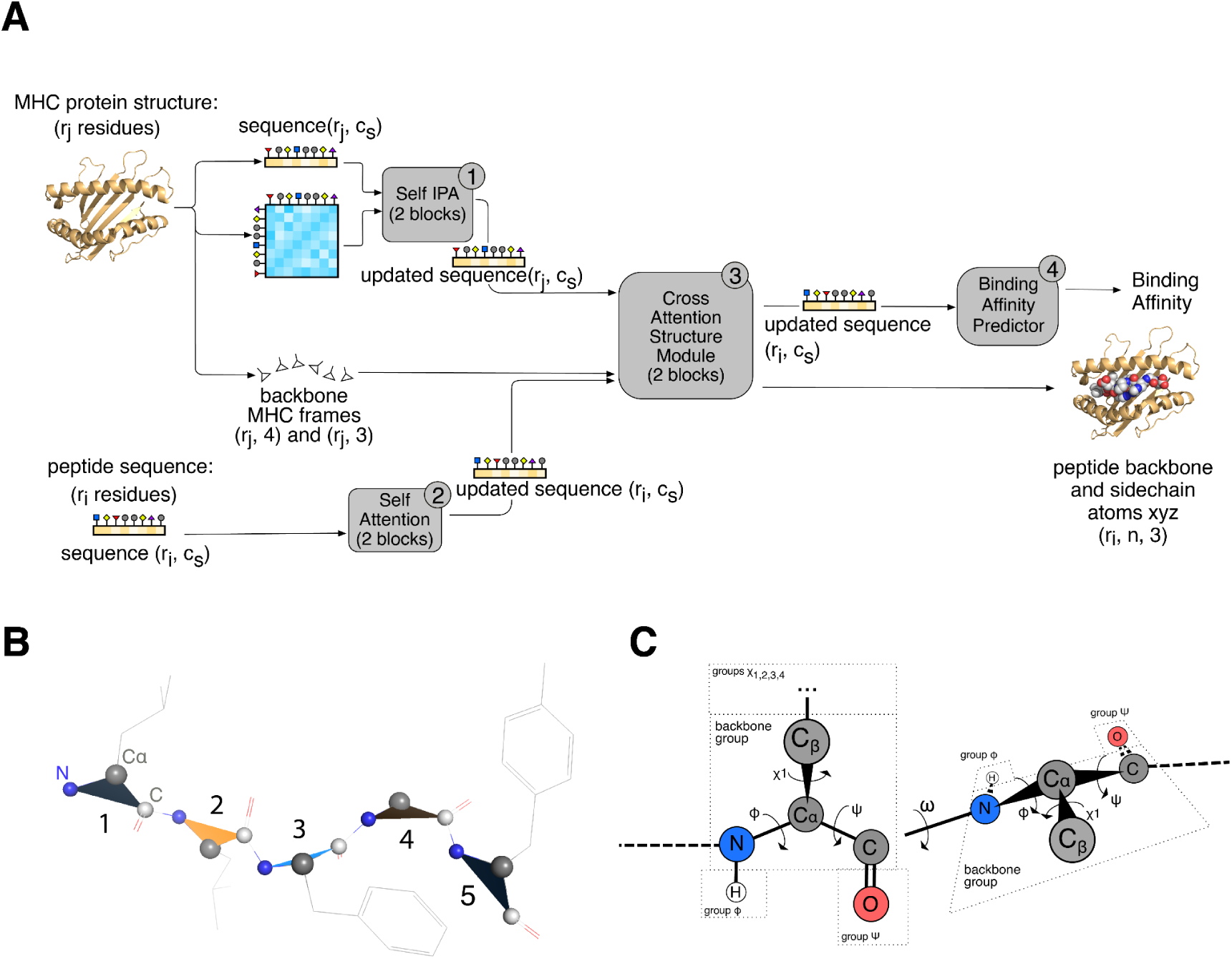
SwiftMHC architecture. **A.** Flow diagram of SwiftMHC. The inputs are the structure of the MHC G-domain, consisting of *r_j_* residues, and a peptide sequence of *r_i_* residues. The model comprises four key modules: (1) the MHC Self Invariant Point Attention (IPA) module, which processes the MHC structure; (2) the Peptide Self Attention module, which processes the peptide sequence; (3) the Cross Attention Structure module, which captures interactions between MHC and peptide residues; and (4) the BA Predictor module. Residue distances in the MHC structure are encoded in a proximity matrix, calculated as 1 / (1 + *d_ij_*), where *d_ij_* is the shortest distance between heavy atoms of residue pairs. The MHC backbone is described using local frames. Both MHC and peptide sequences are represented as one-hot encoded tensors with dimensionality *c_s_* and the structures have maximally *n* atoms per residue. *(c_s_=32 and n=14 in this study)*. **B.** Local frames for a peptide composed of five residues. Backbone local frames are sequentially numbered. Triangles mark the atoms involved in determining the frame orientation (N, Cα and C). **C.** The torsion angle representation of a peptide with two residues. Different peptide conformations can be derived by rotating torsion angles. Each torsion angle corresponds to a rigid group. Every residue has a backbone rigid group and several sidechain rigid groups The atoms in the rigid group are repositioned as the torsion angle is modified. The backbone torsion angles are φ, ᴪ and ω. The ω torsion angles have an empty rigid group, but the rigid groups for φ and ᴪ both contain one atom, though hydrogens are actually only added in the end and only if OpenMM is used. The side chain torsion angles are 𝜒_1_, 𝜒_2_, 𝜒_3_ and 𝜒_4_. Their corresponding rigid groups contain side chain atoms.

With SwiftMHC, we demonstrate the feasibility of designing task-specific AI models that can perform on par with large general-purpose AI systems such as AlphaFold, while being substantially faster. Importantly, SwiftMHC provides both the binding prediction and 3D structural models, crucial for downstream applications such as TCR design. Focusing on HLA-A*02:01 9-mers—given its clinical relevance, prevalence, and the availability of data—provides a robust proof-of-concept. This work highlights the promise of building specialized deep learning (DL) systems on physics-derived 3D models, especially in data-scarce domains such as CDR3 loop modeling for T-cell receptors (TCRs) and antibody-antigen interactions, where evolutionary information is absent. Finally, we outline the current limitations of SwiftMHC and potential avenues for future improvement.

Key contributions of this work include:

1. Showcasing the power of task-specific small AI models trained on physics-derived synthetic data in a data scarce domain, achieving remarkable speed and precision over general-purpose big AI systems such as AlphaFold.
2. Establishing SwiftMHC’s performance for HLA-A*02:01 9-mer peptides, one of the most prevalent alleles in the human population^22–24^, enabling large-scale screening of patient tumor genomes and expanding the repertoire of candidate target peptides.
3. Overcoming the speed bottleneck of structure-based MHC-peptide AI predictors. While structure-based predictors are shown to have better generalizability than the popular sequence-base approaches, their limited speed has hindered scalability.^25,26^
4. Enabling 3D model prediction for safer therapies. By predicting peptide-MHC 3D structures in milliseconds, SwiftMHC can help identify tumor-specific peptides distinct from self-peptides, paving the way for safer and more precise cancer immunotherapies.

## Results

### SwiftMHC architecture

The SwiftMHC network utilizes a residue-to-residue attention mechanism to learn to predict BA and structural features based on interactions between neighboring amino acids. (**Fig. 2A**). Specifically, we apply Self Attention networks on MHC structures and peptide sequences, respectively, to update each residue’s representations with its neighborhood information. Then we apply cross-attention networks to learn the attention weights between MHC and peptide residues, which are expected to encode the interaction information between the peptide and the MHC. This information is used to predict the 3D models and the BA.

Modelling flexible peptides requires efficient representation of residues and their movement in space. The bond angle between nitrogen (N), alpha carbon (Cα), and carbonyl carbon (C) is relatively fixed due to the constraints imposed by the molecular geometry and electronic structure of these atoms. Following AlphaFold2’s approach^15^, we model each residue backbone as a rigid-body triangle with the bond angle N-Cα-C of 109°, allowing free rotation and translation in space while reducing the sampling space (**Fig. 2B**). We use unit quaternions to represent the rotations of a peptide residue as they are more efficient and compact than 3×3 rotation matrices (requiring only 4 parameters vs. 9), and offer better numerical stability. Thus the movement of each residue backbone is modeled as a local frame, which encodes the rotation (i.e., a unit quaternion represented as a vector of 4 ⨉ 1) and translation (a 3 ⨉ 1 vector) of each residue relative to a global coordinate system (**Suppl. Mat. Subsection 1.2.1**). Additionally, the peptide bond torsion angle (ω) is typically 180° or 0°, adding rigidity to the backbone (**Fig. 2C**). We take advantage of this with an auxiliary loss on ω along with other structure losses (**Suppl. Mat. Subsection 2.4.3**). See Methods “Computational Efficiency” for additional design choices improving computational efficiency.

The algorithm is structured into four primary modules (**Fig. 2A**).

1. **MHC Self Invariant Point Attention (IPA) Module** (**Algorithm 2** in **Suppl. mat.**): The purpose of this module is to encode each MHC residue based on its amino acid type and its structural neighbors. This module iteratively applies Self Attention to the MHC structure, drawing inspiration from the AlphaFold2 invariant point attention (IPA) technique to make the residues update each other. It was designed to represent and process interactions between amino acids. For attention weight calculation of each pair of the *r_j_* MHC residues, it utilizes the one-hot-encoded sequence representation of the pairing MHC amino acid types and a *r_j_ x r_j_ x 1* proximity matrix that captures their geometric distances. These attention weights are used for updating the MHC residue features. Those *r_j_*updated feature vectors are the output of this Self Attention submodule.
2. **Peptide Self Attention Module (Algorithm 3** in **Suppl. mat.)**: The purpose of this module is to encode each peptide residue based on its amino acid type and its sequence neighbors. It applies Self Attention iteratively (twice, can be configured) to the peptide sequence, allowing the *r_i_* residues to be notified of each other by means of residue-to-residue updating. Relative positional encoding was used to provide residue positions to the attention network . Updating allows residues to know their position in the peptide, allowing the network to distinguish between them. This peptide Self Attention submodule outputs a sequence of *r_i_*updated peptide residue feature vectors.
3. **Cross Attention Structure Module (Algorithm 4** in **Suppl. mat.)**: The purpose of this module is to update each peptide residue based on the interaction between it and the MHC residues. The updated peptide encoding is used to predict peptide structure and as input of the BA prediction module. This module iteratively performs cross-attention between the *r_i_* peptide residues and *r_j_*MHC residues, employing a Cross IPA on the previously updated residue feature vectors and their geometric distances. Throughout these iterations, the module not only updates the peptide features but also refines the peptide’s structural representation from an initial starting point at the center of the MHC groove. The MHC features are kept fixed in this module. The first step in this structural refinement process is prediction of backbone frames (residue positions + orientations), followed by side-chain torsion angle prediction and finally placement of atoms in the cartesian space.
4. **BA Prediction Module (Algorithm 8** in **Suppl. mat.)**: To allow SwiftMHC to accommodate different peptide lengths, we designed a *residue-wise* BA predictor. The contribution of each residue to BA is predicted by a multi-layer perceptron (MLP), which processes updated peptide residue features produced by the structural module. The end result of the BA predictor is the sum of the MLP outputs over all peptide residues. This BA value is trained to approach 1 - log50000(Kd) or 1 - log50000(IC50), where K_d_ and IC_50_ are experimentally determined binding constants.

The SwiftMHC network is trained by a loss function with multiple terms on BA, FAPE, torsion angle and other structural violations. (**Suppl. Subsection 2.6**). The *BA loss* quantifies the difference between the predicted and true binding affinities between a peptide and the MHC. *The Frame Aligned Point Error (FAPE)* measures the discrepancy in the positions of peptide atoms (both backbone and side chain) between the true and predicted structures. The *Torsion angle loss* assesses differences in torsion angles between the true and predicted structures. *Structural violations*, which include abnormal bond lengths, bond angles, or atomic clashes, are incorporated only during the fine-tuning phase of training (see Methods and **Suppl. Subsection 3.1**).

By default, *SwiftMHC* predicts BA and 3D models without running OpenMM refinement. For BA predictions, users may skip writing predicted 3D models to disk to avoid performance slowdowns, referred to as *SwiftMHC-BA*. Alternatively, enabling short OpenMM refinement produces high-quality models, referred to as *SwiftMHC-OpenMM*.

### SwiftMHC archives SOTA-level BA prediction at sequence-based speeds

In this study we focus on HLA-A*02:01, one of the most prevalent MHC alleles, observed in 95.7% of Caucasians and 94.3% of Native Americans^22^, and has been extensively studied in clinical trials^3^. Additionally, we focused on 9-mer peptides, the most frequent binding peptide length for MHC class I (MHC-I), as shown in **Suppl. Fig. 1A**.

We assessed the BA prediction performance of SwiftMHC using 7,726 BA data points obtained from IEDB^27^. To objectively evaluate SwiftMHC’s performance on unseen peptides, we clustered the data into ten groups and conducted leave-one-cluster-out cross-validation (see Methods). Additionally, we compared SwiftMHC with three state-of-the-art prediction methods: NetMHCpan 4.1^8^, MHCflurry 2.0^7^, AlphaFold2-FineTune^16^ and MHCfold^28^.

To ensure objective evaluation, MHCflurry 2.0 was retrained on the same datasets as SwiftMHC. We note that AlphaFold2-FineTune and MHCfold were not retrained and were evaluated using publicly available pretrained models due to computational restraints. As a result, at least some, if not all, of our test cases were included in the training data of these two methods (see **Methods** for details about the data overlap), which could lead to inflated performance results.

Our results demonstrate that SwiftMHC reliably predicts MHC-binding peptides, achieving a median of AUC of 0.91, outperforming three SOTA methods: AlphaFold2-FineTune, MHCfold and retrained MHCflurry 2.0 (**Fig. 3A and Suppl. Fig. 10AB**). Beyond AUC, SwiftMHC also achieves superior results in terms of area under the precision–recall curve (AUPR) and Pearson correlation (**Suppl. Fig. 10AB**).

**Figure 3.**
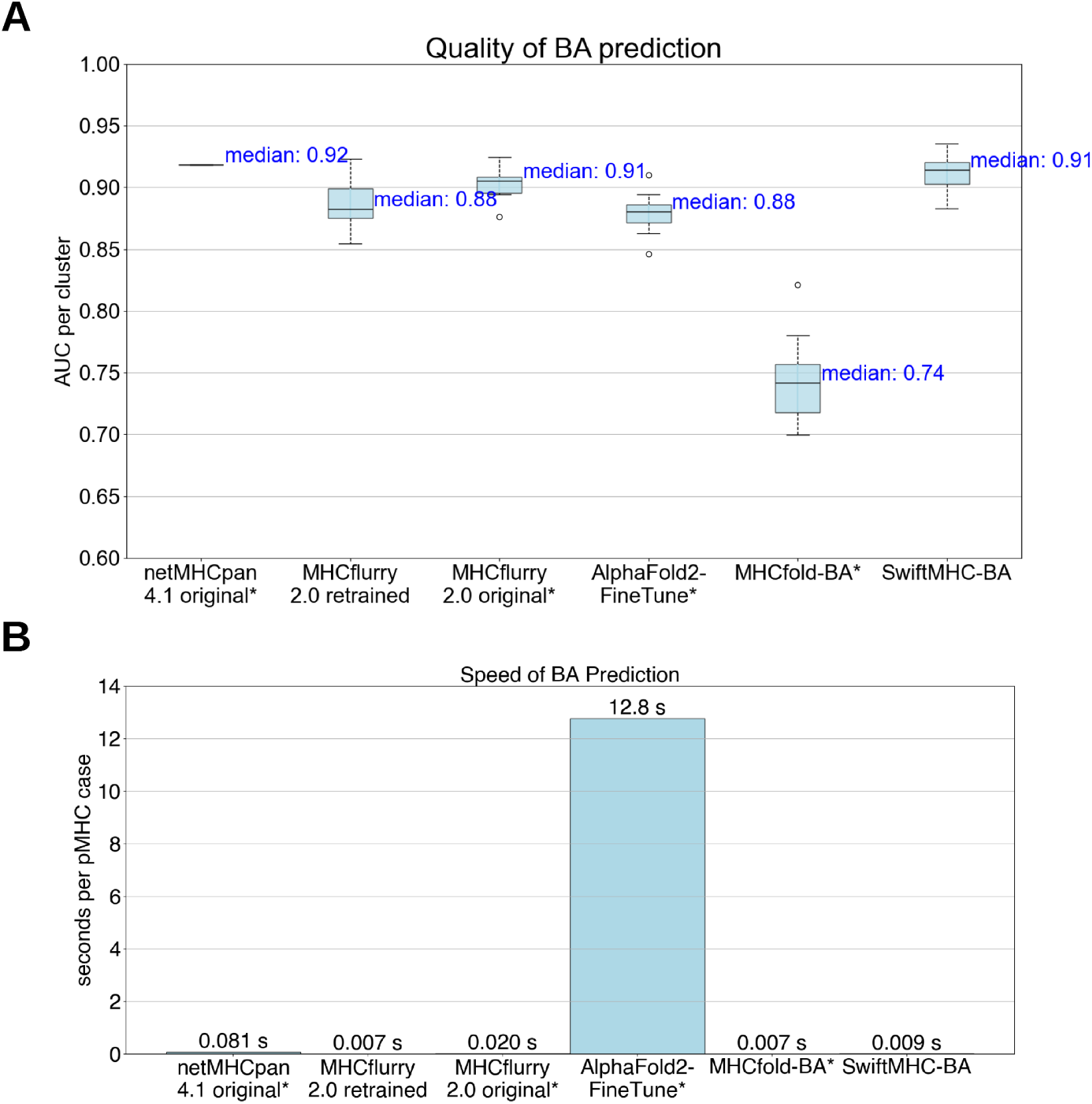
BA prediction quality and speed comparison to SOTA methods. The 7,726 BA data points were separated into 10 clusters. MHCflurry 2.0 was retrained on the same data as SwiftMHC. The asterisks (*) denote the methods that were not retrained and have seen the test data, potentially leading to inflated performance estimates.. **A.** The box plots display the distribution of the AUC for each of the 10 HLA-A*02:01 9-mer peptide clusters. A perfect model has an AUC of 1, while a model with no discrimination has an AUC of 0.5. The box represents the interquartile range (IQR), which is the range between the first quartile (Q1) and the third quartile (Q3). The line inside the box indicates the median (Q2) of the dataset. The whiskers extend from the box to the smallest and largest values within 1.5 times the IQR from Q1 and Q3, respectively. Data points outside of this range are considered outliers and are marked as circles individually. **B.** The bar plot displays the speed of each of the six methods.

Compared to **NetMHCpan 4.1**, SwiftMHC achieves nearly comparable accuracy: NetMHCpan 4.1 attains a median AUC of 0.92, a Pearson correlation of 0.81, and an AUPR of 0.89, whereas SwiftMHC reaches 0.91, 0.80, and 0.88, respectively. Notably, NetMHCpan 4.1’s reported performance may be slightly inflated, as some of its training data overlaps with our test set.

Where SwiftMHC clearly excels is inference speed and structural modeling. It predicts binding in only 0.009 seconds per case on GPU (when predicted structures are not written to disk), even faster than NetMHCpan 4.1 (0.081 s/case; **Fig. 3B**). While MHCfold-BA is marginally faster (0.007s) by omitting sidechain modeling, it exhibits lower AUC and less accurate 3D models (**Fig. 4A**). In contrast, SwiftMHC not only predicts binding affinity but also generates high-quality 3D pMHC models (see below), a critical feature for downstream T-cell receptor (TCR) design.

**Figure 4.**
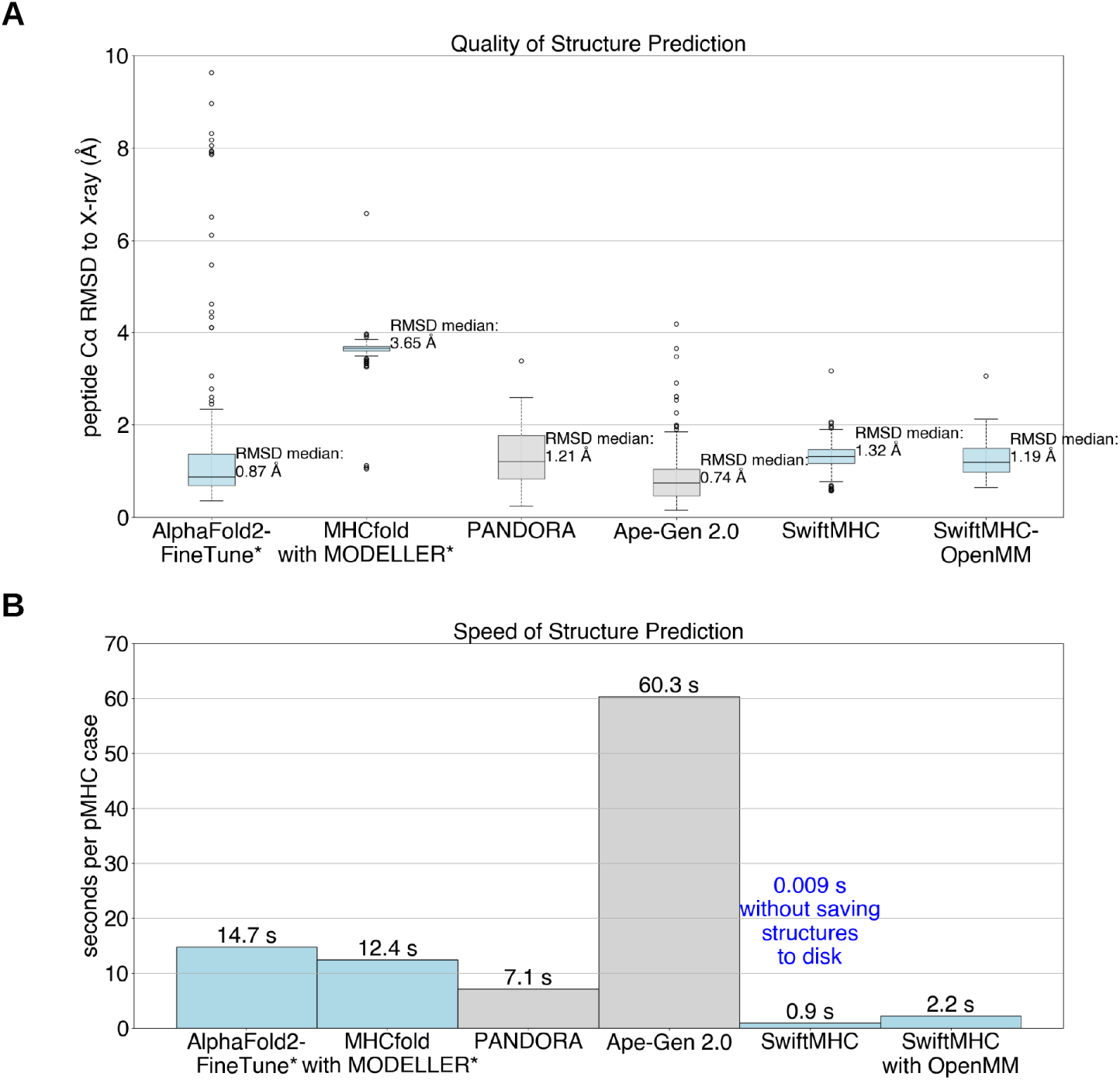
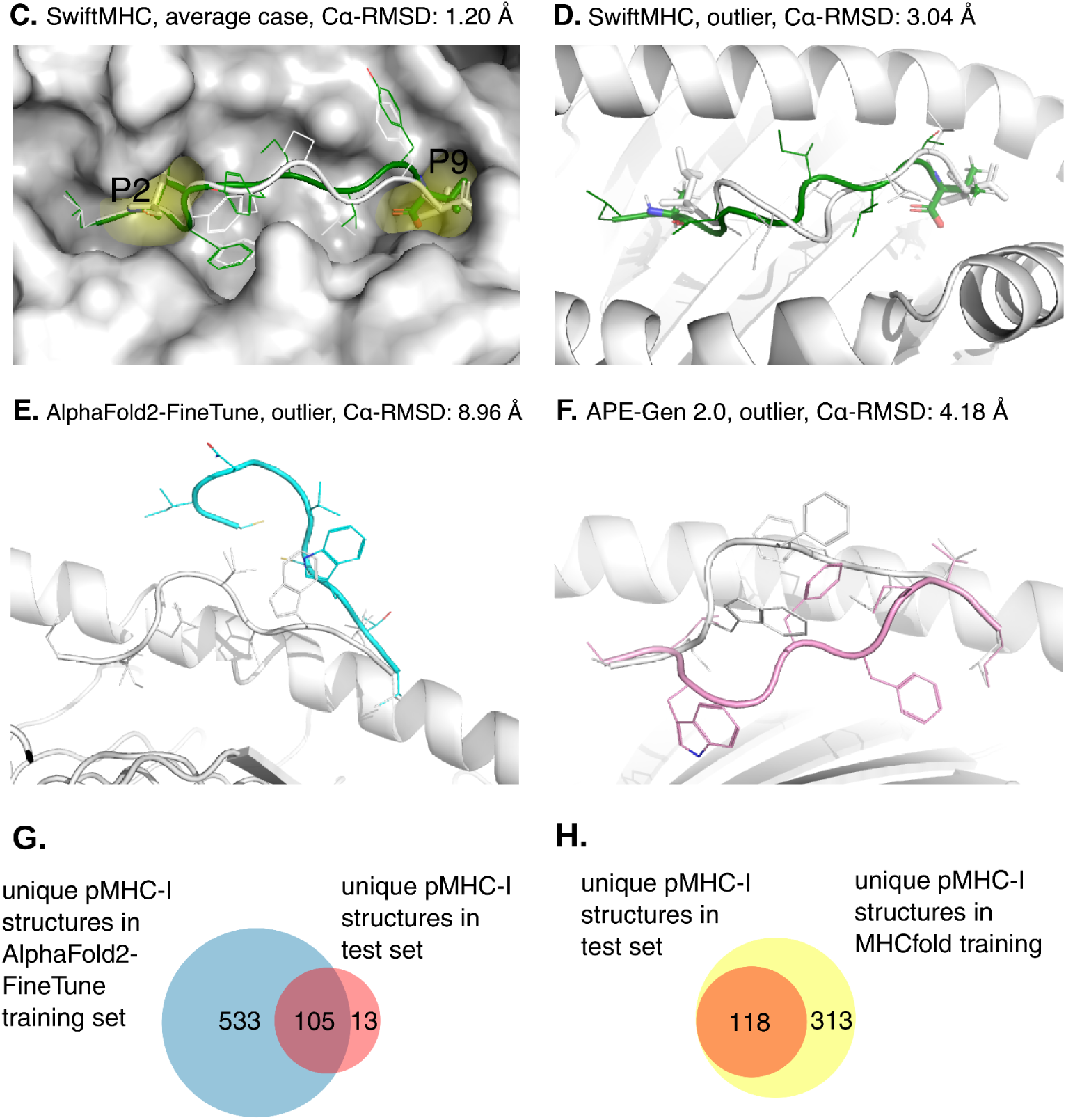
Structural prediction quality and speed assessment on 202 X-ray structures. The 9-mer peptides of 202 HLA-A02:01 X-ray structures were split in 10 clusters and modeled using several methods. AlphaFold2-FineTune and MHCfold are marked with an asterisk (*) as they could not be retrained and may have seen at least part of our test data resulting in inflated results (see Methods and panel **GH** for details about the data overlap). The other methods were employed using a 10-fold cross validation. **A.** Peptide Cα-RMSD values comparing 3D models to X-ray structures. Light blue: DL-based methods. Gray: Physics-based methods. SwiftMHC (with and without OpenMM refinement) performs comparably to SOTA methods. **B.** Average modeling speed per case, measured in batch mode. SwiftMHC achieves 0.01 s/case without outputting structures, processing larger batches from BA data consisting of 7,726 cases total, 533 to 678 cases per cluster, up to 64 cases per batch. With structure output and OpenMM, SwiftMHC achieves 0.9–2.2 s/case reproducing X-ray test data consisting of 202 cases total, 1–57 cases per batch, one batch per cluster. SwiftMHC is the fastest method tested. **C.** Example SwiftMHC model (PDB ID: 1HHK; Cα-RMSD: 1.19 Å), overlaid with its X-ray structure. P2 and P9 anchor residues are labeled. **D.** An exception (PDB ID: 2GTW) where SwiftMHC predicts different anchors than in the X-ray structure, achieving the highest tested Cα-RMSD (3.04 Å). **E.** Outlier by AlphaFold2-FineTune (PDB ID: 3MRG; RMSD: 8.96 Å). N-terminal anchor is incorrect. **F.** Outlier by APE-Gen 2.0 (PDB ID: 2JCC; RMSD: 4.18 Å). The central loop is misaligned, despite correct anchors. **G.** Venn diagram showing the overlap between the unique pMHC-I structures in the training set of AlphaFold2-FineTune and the 202 X-ray structures used as the test set to evaluate the performance of all structure prediction methods in this study. **H.** Venn diagram showing the overlap between the unique pMHC-I structures in the training set of MHCfold and the X-ray structures used as the test set to evaluate the performance of all structure prediction methods in this study.

SwiftMHC accurately predicts both high and low affinity binders. We evaluated SwiftMHC’s performance across different binding affinity ranges by analyzing a confusion matrix (**Suppl. Fig. 10E**). The results show that SwiftMHC predicts most accurately for peptides with high (50–500 nM) and low (5000–50000 nM) binding affinities. This indicates that the model does not systematically overpredict binding and is capable of distinguishing non-binders effectively.

### SwiftMHC is robust to peptide single point mutations

To assess SwiftMHC’s robustness to single-point mutations, we predicted BA for 2,838 single amino acid variants derived from the original dataset of 7,726 peptides using the network models trained on the wild-type cluster’s respective folds and the MHC structure from PDB entry 3MRD. For each mutant, we calculated the true change in ΔG (ΔΔG) relative to its wild-type counterpart and compared this to the predicted ΔΔG (**Suppl. Fig. 10F**). The overall Pearson correlation between predicted and true ΔΔG values is relatively low (r = 0.33). Focusing specifically on mutations that do not shift a peptide from non-binder to binder status (IC_50_/K_d_ < 500 nM) or vice versa, SwiftMHC incorrectly classified only 11% of either the wild-type or the mutant. This indicates that while fine-grained ΔΔG prediction remains challenging, SwiftMHC reliably captures whether a mutation is functionally impactful in terms of binding.

### SwiftMHC delivers Angstrom-level accuracy in all-atom pMHC structure prediction with high computational efficiency

We evaluated SwiftMHC and four SOTA methods for modeling 3D pMHC-I structures: AlphaFold2-FineTune^16^ (DL-based), MHCfold^28^ (DL-based), PANDORA 2.0 (physics-based)^12^ and Ape-Gen 2.0 (physics-based)^13^ (**Table 1**).

**Table 1.**
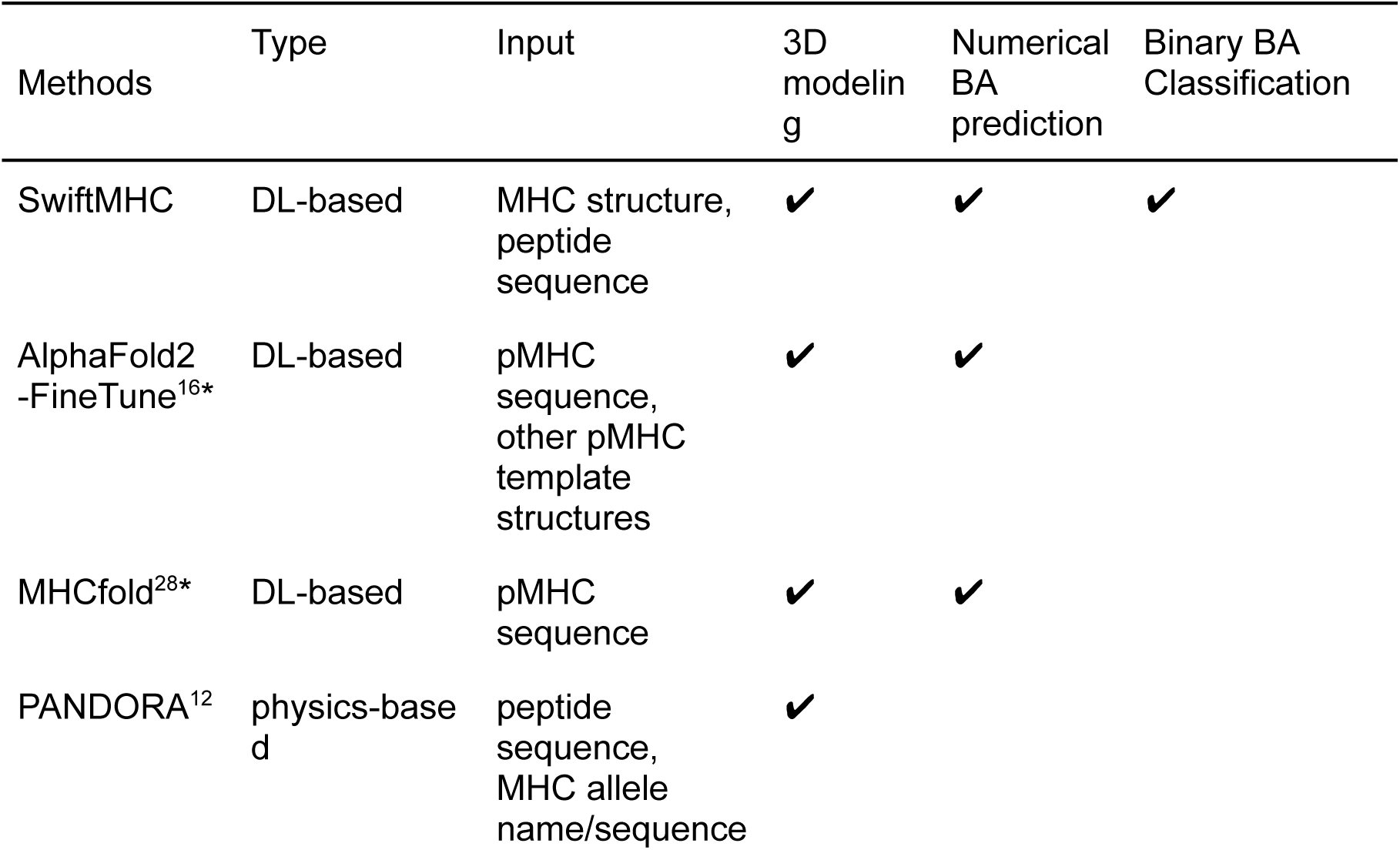
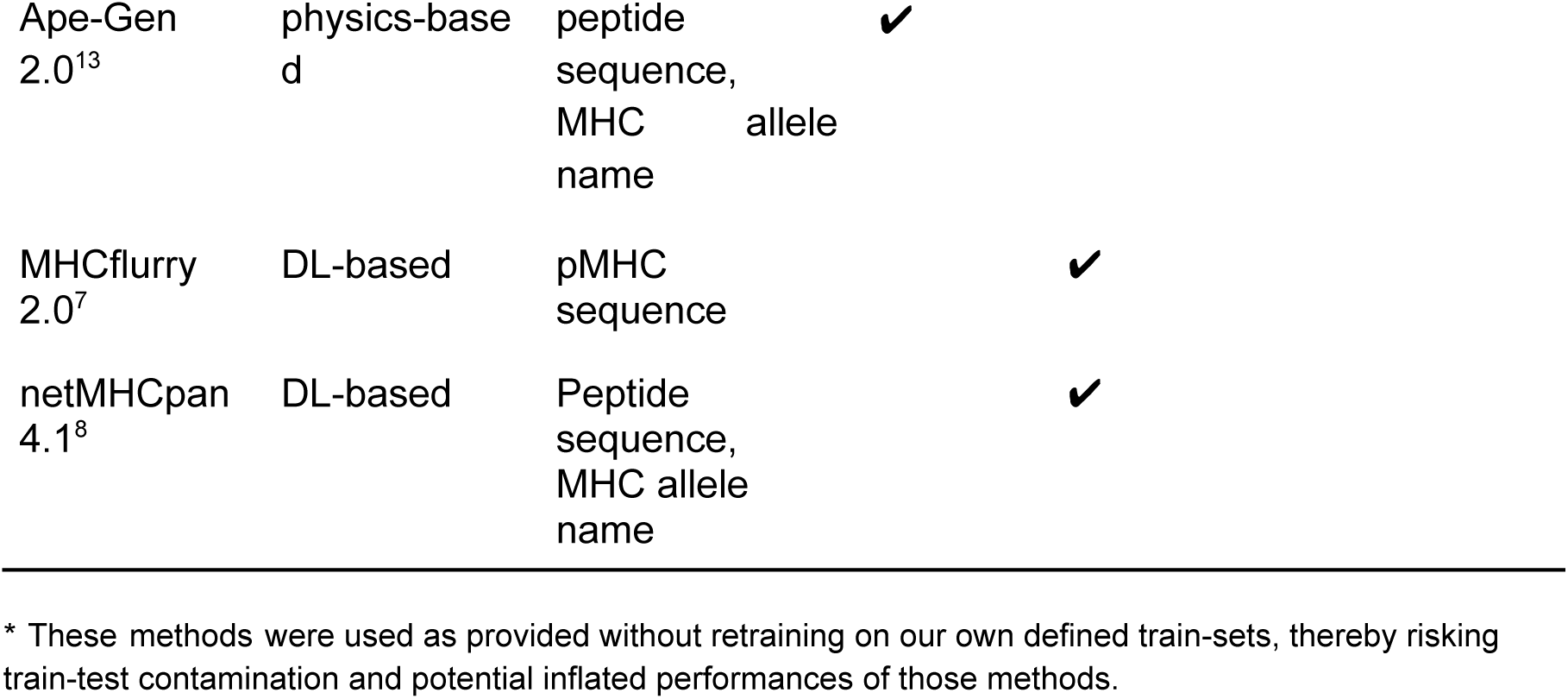
Methods evaluated for BA prediction and structure prediction.

To objectively evaluate the performance, we clustered 202 HLA-A*02:01 9-mer X-ray structures together with the PANDORA models for the 7,726 IEDB entries based on the peptide sequence similarity (**Suppl. Fig. 1B**). We ensured that the X-ray structures and PANDORA 3D models from the same cluster were not included in the training datasets. PANDORA and Ape-Gen 2.0 rely on homology modeling, using X-ray structures as templates to predict new pMHC-I 3D conformations. To ensure unbiased evaluation, we excluded any X-ray structures containing the same peptide to prevent their use as templates by these methods.

SwiftMHC-OpenMM produces highly accurate pMHC-I 3D models with a median Cα-RMSD of 1.19 Å (**Fig. 4A**), comparable to three other state-of-the-art methods (median Cα-RMSD: 0.87–1.21 Å), significantly outperforming MHCfold (3.65 Å). Although AlphaFold-FineTune has a lower median Cα-RMSD of 0.87Å, this advantage likely reflects substantial overlap between its training and our test data. Importantly, SwiftMHC produces far fewer outliers than AlphaFold-FineTune (**Fig. 4)**, highlighting the benefit of augmenting training with task-specific, physics-derived 3D models.

In terms of computational efficiency, SwiftMHC processes each case in 0.9 seconds (0.009 seconds per case when excluding PDB file writing) and 2.2 seconds with OpenMM energy minimization (**Fig. 4B**). When disk writing is bypassed, SwiftMHC is up to thousands of times faster than existing state-of-the-art 3D modelling approaches.

*Outlier analysis.* The peptide-binding groove of HLA-A*02:01 has two primary deep pockets, P2 and P9, which are critical for peptide binding and stabilization^29^. With few exceptions (e.g., 2GTW, discussed below), the P2 and P9 pockets commonly anchor the 2nd and the 9th residues of the peptide. Consistently, most of the 3D models from SwiftMHC have a backbone structure similar to their X-ray counterpart, with common anchor positions 2 and 9 (**Fig. 4C**). The top outlier for SwiftMHC and PANDORA is 2GTW, of which the X-ray structure has anchor positions 1 and 9, but it is predicted with anchor positions 2 and 9 (**Fig. 4D**). Since PANDORA uses netMHCpan 4.1 for anchor prediction, and SwiftMHC was trained on PANDORA models, SwiftMHC likely inherits these netMHCpan-based prediction errors, reducing accuracy in such cases.

MHCfold exhibited a notably higher median Cα-RMSD (3.65Å) compared to the other methods (0.87-1.32Å). Although AlphaFold2-FineTune achieved a low median Cα-RMSD, it produced many outliers, including at least two 3D models in which one end of the peptides was positioned outside the MHC groove (**Fig. 4E**). In contrast, in the top three outlier 3D models generated by Ape-Gen 2.0, the peptide center loop was overly buried within the MHC groove, making it less surface-exposed compared to the corresponding X-ray structures(**Fig. 4F**).

### Structural analyses reveal that SwiftMHC-generated 3D models closely approximate the quality of X-ray structures

In addition to RMSD, we assessed model quality by evaluating van der Waals clashes, chirality, and backbone dihedral angles.

*Number of van der Waals clashes.* The OpenMM energy minimization step in SwiftMHC effectively reduces van der Waals clashes (see definition in Methods), which occur when atoms are positioned too closely. Across the 202 test cases, these reductions are comparable to levels achieved by other methods (**Fig. 5A**), highlighting the importance of a final molecular dynamics step to ensure the physical realism of the 3D models.

**Figure 5.**
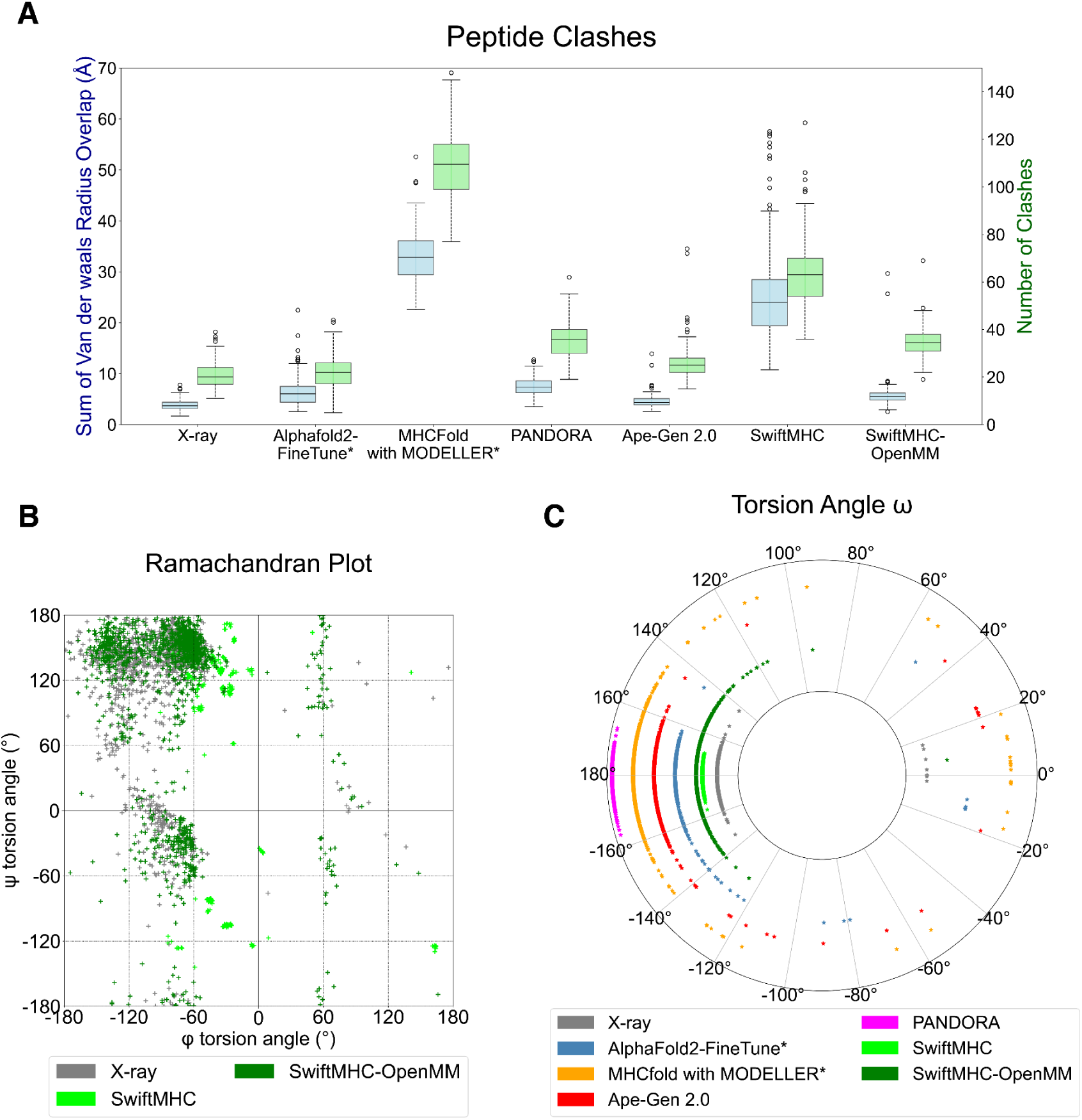
Structural characterizations of SwiftMHC-generated 3D models for the 202 x-ray test cases. **A.** Distribution of number of clashes and the sum of their van der Waals radii overlaps per 3D model/structure. **B.** Distribution of backbone φ and ψ torsion angles as a Ramachandran plot for both SwiftMHC 3D models and the X-ray structures. **C.** Distribution of the ⍵ peptide bond torsion angle for all 3D models and X-ray structures.

*Chirality.* Chirality in amino acids refers to the property where an amino acid molecule is non-superimposable on its mirror image: L- and D-enantiomers. Chirality plays a crucial role in the structure and function of biomolecules, as the spatial arrangement of groups affects how amino acids interact with other molecules. Except for glycine, amino acids are chiral molecules and the vast majority of amino acids in proteins and enzymes of living organisms are in the L-form. In the 202 test x-ray cases, some amino acids lacked side-chain atoms, which we restored using OpenMM PDBfixer^30^. However, in some instances, PDBfixer introduced D-form amino acids. Since both SwiftMHC and AlphaFold2-FineTune use these structures as input for model generation, we specifically examined D-form amino acids that were introduced in the models (**Table 2**). SwiftMHC maintains chirality by rotating and translating local frames of amino acids, as well as placing the chiral hydrogen before executing OpenMM. As a result, it produced L-form amino acids in the generated peptide structures. However, when OpenMM was used for energy minimization, some L-form amino acids were converted to D-form. APE-Gen 2.0, which also relies on OpenMM, exhibited the same issue and even introduced D-form amino acids in the peptide structure. In contrast, AlphaFold2-FineTune and PANDORA, which do not use OpenMM, did not introduce any D-form amino acids. MHCfold, which predicts individual atomic positions around the chiral Cα atom, introduced a D-form amino acid in one of its models.

**Table 2.** Methods evaluated for amino acid chirality in predicted structures.

| Methods | PDBfixed*<br>X-ray<br>structures<br>used as<br>input | OpenM<br>M used | Average<br>number of<br>chiral amino<br>acids per<br>model** | Number of<br>D-amino<br>acids<br>introduced<br>in the MHC | Number of<br>D-amino<br>acids<br>introduced<br>in the<br>peptide |
| --- | --- | --- | --- | --- | --- |
| MHCfold with<br>MODELLER |  |  | 174 | 1 | 0 |
| AlphaFold2-F<br>ineTune | ✓ |  | 173 | 0 | 0 |
| APE-Gen 2.0 |  | ✓ | 263 | 22 | 12 |
| PANDORA |  |  | 263 | 0 | 0 |
| SwiftMHC | ✓ |  | 174 | 0 | 0 |
| SwiftMHC-O<br>penMM | ✓ | ✓ | 174 | 9 | 0 |

\*OpenMM PDBfixer<sup>30</sup> was used to restore missing side-chain atoms in some of the 202 X-ray structures, which resulted in the introduction of 11 D-form amino acids.
\*\*Some methods model the entire class I MHC protein structure, while others model only the G-domain, leading to differences in the total number of amino acids.

*Dihedral angles of the peptide backbone*. The backbone torsion angles, ϕ (phi) and ψ (psi), define the conformation of a protein’s backbone. These angles are constrained by steric clashes, such as repulsions between side-chain atoms or backbone atoms, which limit the range of energetically favorable conformations a protein can adopt. Validating computational 3D protein models includes verifying that the dihedral angles fall within the naturally observed regions of the Ramachandran plot, as deviations can signal errors in the model. To this end, we generated Ramachandran plots for the X-ray structures and the 3D models produced by all six methods (**Fig. 5B** and **Suppl. Fig. 9**). The φ and ᴪ torsion angles produced by the DL part of SwiftMHC are restricted to several small areas in the Ramachandran plot (**Fig. 5B**). After energy minimization in OpenMM these angles were more widespread and the SwiftMHC distribution became more closely with that of experimentally determined X-ray structures.

*ω torsion angles and Cis/trans configurations*. "Trans" and "Cis" refer to the geometric configuration around peptide bonds, specifically involving the arrangement of the atoms adjacent to the peptide bond (the amide bond between the carbonyl carbon of one amino acid and the amide nitrogen of the next). The trans configuration is the most common configuration found in proteins, as it is generally more stable due to less steric hindrance between the side chains of the amino acids. In the trans configuration, the dihedral angle around the peptide bond is approximately 180°.

All five methods generated 3D models in which the ω torsion angles do not correspond to those observed experimentally in the X-ray structures (**Fig. 5C**). For instance, in the case with PDB ID 5SWQ, all methods predict an asparagine-glycine peptide bond with an ω angle ranging from 160.0° to 200.0° (trans), whereas the X-ray structure shows this angle as -0.6° (cis). Similarly, SwiftMHC predicts seven glutamate-proline peptide bonds with ω angles between 160.0° and 200.0° (trans), while the X-ray structures indicate that these angles fall between -20.0° and 20.0° (cis). Notably, OpenMM appears to shift these specific angles outside the trans range, approaching 140°. Furthermore, SwiftMHC predicts nearly all ω angles in the trans-conformation, although the energy minimization step in OpenMM slightly alters these angles in some cases. An extreme case is the isoleucine-proline bond in the SwiftMHC 3D model for 5ENW. This ω angle was almost flipped by OpenMM from -178° (trans) to 7° (cis).

### Combined BA and Structural Training Enhance Predictive Performance

We assessed whether the BA data and structural modeling enhance each other’s predictive performance. For this, we conducted 10-fold cross-validation with either BA or structural loss terms disabled, measuring the performance of the other. SwiftMHC-openMM was used in this experiment.

We found that the predicted 3D models deviated significantly more from their corresponding X-ray structures when BA losses were excluded from training (**Fig. 6A**). Many models showed errors, particularly in modeling peptide anchor positions, where this did not happen when BA loss was included in the training (**Fig. 6C**). Additionally, when SwiftMHC was trained without structural loss terms (i.e.,FAPE, torsion, bond length violations, bond angle violations and clashes), the quality of BA predictions was substantially reduced, with the median AUC dropping from 0.91 to 0.72 (**Fig. 6B**). These findings demonstrate that the experimentally determined BA data and the predicted structural models complement each other, enhancing their combined predictive performance.

**Figure 6.**
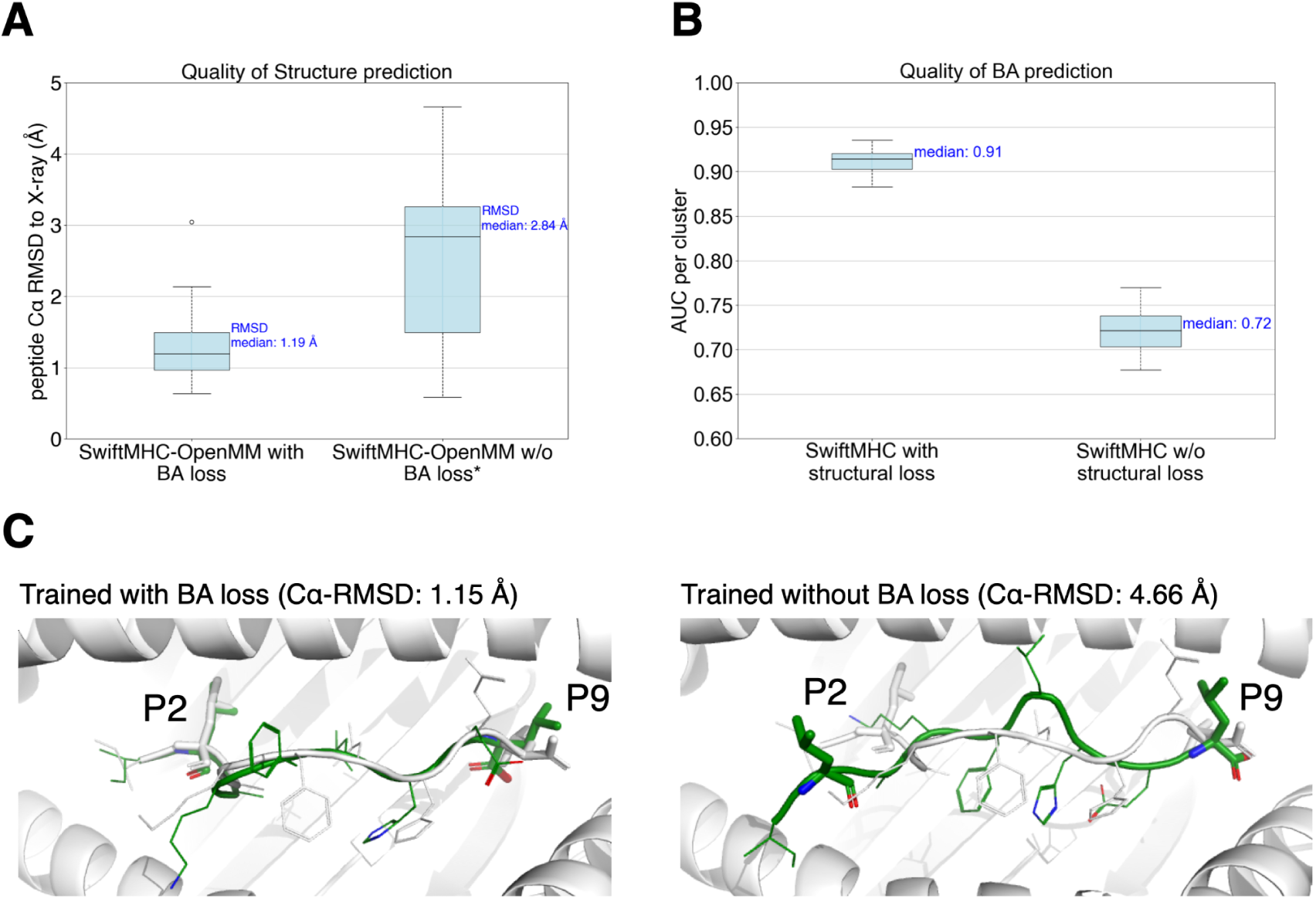
The impact of structural and BA loss terms on structural or BA prediction performance. **A.** The quality of SwiftMHC-generated OpenMM-refined 3D models for regular training vs. training without BA loss, illustrating the BA data enhanced the 3D model quality. *Note that the 3D model for PDB entry 1BD2 is missing from the set of 3D models created without BA loss, as this particular model was structurally compromised to the extent that OpenMM could not successfully perform energy minimization. **B.** BA prediction performance for regular training vs. training without structural losses, showing PANDORA models enhanced the accuracy of BA prediction. **C.** An example case (PDB ID: 5MER) to show the models generated by regular training (left) vs. training w/o the BA data (right) . Green: SwiftMHC 3D models refined by OpenMM; White: X-ray structures. Without the BA loss in the training, SwiftMHC failed to generate correct anchors for the peptide (right panel, peptide Cα-RMSD: 4.66 Å), while with regular training it can generate realistic peptide conformation (left panel, peptide Cα-RMSD: 1.15 Å). Anchor positions 2 (P2) and 9 (P9) are labeled.

## Discussion

We introduce SwiftMHC, an attention-based neural network for the rapid and accurate identification of MHC-bound peptides and generating 3D structures. SwiftMHC delivers high-resolution *all-atom* 3D models with a median Cα-RMSD of 1.19 Å, with a remarkable efficiency by processing each case in just 0.009 (AI mode) to 2.2 (AI+openMM mode) seconds in batch mode on a single A100 GPU. SwiftMHC effectively overcomes the speed bottleneck of structure-based BA predictors while maintaining exceptional accuracy in both binding affinity prediction and 3D structure modeling.

The speed and accuracy of SwiftMHC is attributed to two key innovations: 1) removing the computationally demanding MSA attention module—since MHC structures are conserved and peptides lack evolutionary information—and 2) incorporating physics-derived pMHC-I 3D models into training. Whereas the AI industry often focuses on large, general-purpose models like AlphaFold313, these approaches demand vast datasets, extensive compute resources and high energy consumption, both during training and inference, while remaining difficult to optimize In contrast, SwiftMHC demonstrates that smaller, task-specific networks – when augmented with extensive physics-based synthetic data – can achieve superior efficiency without sacrificing accuracy. This design highlights a sustainable path forward for domain-focused biomolecular modelling.

In addition, we explored whether cross-attention weights could provide interpretable signals for prediction confidence. By mapping attention scores onto the MHC surface (**Suppl. Fig. 11C–D**), we observed distinct patterns between correct (1HHK, anchors at positions 2 and 9) and incorrect (2GTW, anchors at positions 1 and 9 but predicted as 2 and 9) anchor assignments. In the correct case, the model concentrated attention on the P2 pocket, while in the mispredicted case such localized focus was absent. This suggests that attention distributions may serve as indicators of anchor misprediction, although systematic validation will be required to establish their utility.

As the fastest pMHC-I structure predictor, SwiftMHC facilitates large-scale peptide screening for vaccine development and immunotherapy research. Its high efficiency enhances the ability to identify novel neoantigens from the diverse range of tumor-derived peptide fragments, while providing detailed 3D models for studying molecular interactions and T-cell recognition. Furthermore, SwiftMHC’s speed offers the potential to construct the first comprehensive 3D self-peptide-HLA (human MHC) library. Such a library could be pivotal for identifying TCR-therapy peptide targets that differ from self-peptides on T-cell-exposed surfaces, ultimately contributing to the development of safer and more effective therapeutic strategies.

This proof-of-concept study focuses on HLA-A*02:01 9-mers, selected for their prevalence and clinical relevance. Their abundant data and fixed peptide length makes it a relatively straightforward case for sequence-based predictors used in comparison. A key direction for future work is to expand SwiftMHC training set to include additional alleles and peptides of varying lengths. Such expansion is expected to enhance SwiftMHC’s generalizability, aligning with the reported 6–16% performance gains of structure-based methods over sequence-based methods^25^.

Lastly, while optimized for pMHC complexes, our SwiftMHC strategy can be adapted for broader applications, including protein-peptide interactions and CDR3 loop modeling for antibody-antigen and TCR:pMHC-I complexes. The flexibility and the lack of evolutionary signal of peptides and CDR3 loops pose significant challenges for accurate modeling, even for advanced methods like AlphaFold^31^. Capturing peptide and CDR3 conformational changes upon binding is critical for advancing therapeutic TCR and antibody design and improving our understanding of immune recognition and defense.

### Model Limitations

This proof-of-concept study has several limitations. First, SwiftMHC is currently restricted to HLA-A*02:01 9-mer peptides. However, its algorithm is designed to easily incorporate training sets for additional alleles and peptides of varying lengths.

Second, for computational efficiency, we choose to design SwiftMHC to use fixed MHC orientations not allowing random orientation options. This limitation could be addressed by redesigning the network to incorporate rotation-translation equivariance, such as using backbone frames based on the MHC protein’s intrinsic orientation rather than identity frames.

Third, SwiftMHC does not yet account for peptides with post-translational modifications (PTMs) such as phosphorylation, glycosylation, methylation, and acetylation. Addressing this would require training on datasets with sufficient examples of PTM-modified structures. While SwiftMHC cannot currently model PTMs directly, it can generate reasonable starting conformations for downstream molecular dynamics (MD) simulations of modified peptides.

Finally, the speed evaluation of our study was conducted using a maximum batch size of 64 pMHC cases, optimized for large datasets (∼8000 cases, as in the BA test datasets) that efficiently filled most batches. However, SwiftMHC’s efficiency may decline when processing very small datasets with only a handful of cases per batch. Additionally, the impact of software initialization time, which may reduce efficiency for smaller datasets, was not explored.Further investigation into time consumption for larger datasets could offer additional insights into its performance efficiency.

## Methods

### Data composition

In this study, we focus on HLA-A*02:01 alleles and 9-mer peptides. For this, 7,726 BA data points were collected from IEDB^27^ and 202 X-ray structures were collected from the PDB^10^. OpenMM PDBfixer^30^ was used to fill in missing heavy atoms in these X-ray structures in cases where a residue was incomplete. To critically evaluate the generalizability of our predictor and to prevent data leakage, BA and X-ray data were merged together and Gibbs clustered^32^ based on their peptide sequences. This was performed using GibbsCluster 2.0, with a cluster similarity penalty (λ) set to 0.8 and a small cluster weight (σ) set to 5. The data points were separated into 10 clusters, each of which was used for testing SwiftMHC using a leave-one-cluster-out cross-validation principle (**Suppl. Fig. 1B**).

From the left-out cluster of each fold, the BA data points were used to evaluate the BA prediction quality, while the X-ray structures from that same cluster were used for evaluating structure prediction quality. The BA data points of the remaining 9 clusters were used for training (90% of data) and validation (10%). X-ray crystallographic data were excluded from the training and validation datasets to assess the network’s 3D modeling performance when trained solely on physics-based 3D model data, though including X-ray data would likely enhance our performance.

Since SwiftMHC requires structural data for training, we generated 3D models for each pMHC-I complex in the BA dataset using PANDORA. For each pMHC-I complex, the 3D model with the lowest molecular probability density function score (molpdf) energy score was selected to be used as input to SwiftMHC.

### Data Preprocessing

SwiftMHC gradually updates the peptide structure from identity frames (**Suppl. Algorithm 4**), that is, the starting positions of all peptide residues are at the origin and their N-Cα-C plane overlaps so that they all have the same orientation. This design requires all input structural data in the same orientation, so we superimposed the pMHC-I complex of all X-ray structures and PANDORA 3D models on a reference MHC structure (PDB ID: 3MRD) before training and testing. The reference structure was prepared by isolating the G-domain and placing the geometric center of the MHC binding groove at the origin. All other structures or PANDORA 3D models were structurally aligned to this reference using the PyMOL align command^33^ with its default settings: 2.0 Å as outlier rejection cutoff, 5 outlier rejection cycles and using BLOSUM62 for sequence alignment.

Additionally, some values were pre-calculated from the aligned pMHC-I structures, so that SwiftMHC can quickly access them. Those variables were: *ground truth frames*, *ground truth torsion angles*, *MHC proximity matrix*, *peptide and MHC amino acid sequences*, *residue masks* and *ground truth BA*. See **Suppl. Subsection 1.2** for details about these variables.

### Software and libraries used

SwiftMHC was developed using Python 3.12 (www.python.org) and implemented with the PyTorch 2.0.1 library^34^ for deep learning functionalities. OpenFold 1.0.0^35^ was utilized for key tasks such as FAPE and torsion loss calculations, deriving frames from atomic coordinates, reconstructing atomic coordinates from predicted frames, and performing frame operations. The design of SwiftMHC’s IPA modules was also partially inspired by OpenFold code. For handling PDB files, BioPython 1.8.4^36^ was employed for parsing and representation. Structural refinements were performed using OpenMM 8.1.1^30^.

### Training

Ten SwiftMHC network models were trained for each of the 10 peptide clusters, using 10 distinct training datasets. Training consisted of two phases (**Suppl. Subsection 3.1**): the first phase takes a maximum of 1000 epochs and back propagates on loss terms, based on BA regression and amino acid similarity between true and predicted peptide structure. The second phase takes a maximum of 100 epochs and includes additional loss terms, based on bond length, bond angle and interatomic distance violations (**Suppl. Subsection 2.6**).

During training, both phases were executed for each network model. An early stopping condition was applied, allowing a phase to terminate early if the total loss on the validation dataset ceased to improve (**Suppl. Section 3.2**). The network model with the lowest total loss on the validation dataset in the first phase was selected to be the initial model in the next phase.

### Post Processing the Resulting Structures

Once a pMHC-I structure is predicted and PDB to disk writing is enabled, SwiftMHC optionally runs OpenMM^30^ to minimize its energy using an amber99sb force field. To prevent chirality issues in non-glycine backbones, the hydrogen atom attached to the chiral Cα atom is added by SwiftMHC before running OpenMM. All other hydrogen atoms are added by OpenMM.

### Computational Efficiency

For speed purposes, several design choices were made for SwiftMHC. Since the MHC structure remains fixed, its backbone frames and distance matrix can be precalculated and reused throughout the prediction process. This avoids repeated computations. The preprocessed data was stored in an HDF5 file format to facilitate quick access and efficient data management. For an efficient cross attention computation, MHC residues that do not lie close to the peptide are masked out, meaning that they do not contribute to any operation in that module (**Suppl. Fig. 3**). PDB structures are generally only written to disk on demand and never during training. In addition, we replaced the traditional 3x3 rotation matrix operations with quaternion operations, reducing numerical computations and improving efficiency.

The *time consumption* of executing a fully trained SwiftMHC network model was measured on a single NVIDIA A100 GPU (40 GiB HBM2 memory with 5 active memory stacks per GPU) and 18 Intel Xeon Platinum 8360Y CPUs and 40 GiB 3.2 GHz DDR4 memory available. We made SwiftMHC allocate 16 worker processes to read input data and 16 builder processes to write out structural data in PDB format. For comparing speed, SOTA methods were executed on the same hardware configuration as SwiftMHC. We measured the exact time duration of every method to complete with one-hundredth of a second precision.

### SOTA methods

We compared SwiftMHC against several SOTA methods (**Table 1**) on BA prediction and/or structure prediction qualities. Five methods were compared for their speed and BA prediction quality: SwiftMHC, AlphaFold2-FineTune^16^, MHCflurry 2.0^7^ (both original and retrained models), netMHCpan 4.1 and MHCfold^28^. The AUC was used as a BA quality metric. We classified a peptide as binding to its MHC if IC_50_ or K_d_ was below 500 nM and as non-binding otherwise. Six methods were compared for reproducing the 202 X-ray structures: SwiftMHC, SwiftMHC-OpenMM, AlphaFold2-FineTune^16^, PANDORA^12^, APE-Gen 2.0^13^ and MHCfold^28^. For comparing BA prediction or structure generation speed, all SOTA methods were executed on the same hardware configuration as SwiftMHC.

*AlphaFold2-FineTune* is a modified version of AlphaFold2, where the weights and biases of the network model have been fine-tuned using pMHC BA and X-ray pMHC structural data. AlphaFold2-FineTune achieves faster processing times than standard

AlphaFold2 by omitting template searching and multiple sequence alignments. Instead, it requires the user to provide a query-to-template alignment. A second reason for the increased speed of AlphaFold2-FineTune compared to the standard AlphaFold2 tool is that it does not refine its output structures using an Amber forcefield in OpenMM, which is typically employed for energy minimization in the standard AlphaFold2 pipeline.

We used the 202 X-ray HLA-A*02:01 9-mer structures as templates for AlphaFold2-FineTune. For selecting templates, the Gibbs clustering was used. This means that for each target peptide from a specific cluster, the pMHC-I X-ray structures from the other nine clusters were selected as the template set. To create MHC alignments as input for AlphaFold2-FineTune, the target G-domain MHC sequences were aligned to the template MHC sequences using the BioPython pairwise alignment tool^36^ with default settings (Needleman-Wunsch algorithm, match score 1, mismatch score 0, gap scores 0). For peptide alignment, the 9-mer peptides were aligned in a one-to-one manner (i.e., position one aligned with position one, position two with position two, and so on). Other components of the X-ray structures, such as β2-microglobulin and the ɑ3 domain, were excluded from the alignments. The time to make these alignments was not included in the computation time of AlphaFold2-FineTune. The quality of structural predictions was assessed by comparing the 202 X-ray HLA-A*02:01 9-mer structures against 3D models predicted by AlphaFold2-FineTune. The total runtime required to process these 202 entries was documented.

To assess the quality of BA predictions, we compared 7,726 experimental BA values against predictions generated by AlphaFold2-FineTune. Due to redundancy in the pMHC-I sequences (sometimes there are multiple BA values per complex) and the computational demands of running AlphaFold2-FineTune, we ran AlphaFold2-FineTune on the 6,057 non-redundant data. The total runtime required to process these 6,057 entries was documented. This evaluation aimed to measure AlphaFold2-FineTune’s ability to differentiate true binders (experimental K_d_ or IC_50_ < 500 nM) from true non-binders, following a similar approach to that described in ^16^. For each pMHC-I entry, the negative mean of the MHC-to-peptide and peptide-to-MHC position-aligned error (PAE) values was used as a predictive score. These scores were compared with experimental BA values to calculate the AUC for each cluster. The prediction scores that correspond to the repeated sequence entries were replicated, ensuring each experimental BA value was compared to its corresponding prediction score.

*MHCfold*^28^ is a deep learning tool that first predicts the structures of both the peptide and the MHC molecule from their sequences using convolutional neural networks (CNNs). It then employs a modified version of multi-headed attention along with a structure-derived distance matrix, similar to IPA, to predict BA as either binding or non-binding. To evaluate the BA and structural prediction quality of the MHCfold algorithm, we used the MHC G-domain sequences along with corresponding peptide sequences as inputs. MHCfold’s reliance on side chain modeling and PDB generation may slow down processing. For fair comparison with SwiftMHC-BA, these features were disabled during BA evaluation (referred to as “MHCfold-BA”). We used MHCfold to predict BA for all 7,726 data points and calculated the AUC based on the predictions. The total runtime was measured. Additionally, MHCfold was used to generate pMHC structures for all 202 X-ray complexes. For this. MHCfold was set to predict both the 3D structure and BA and to use MODELLER for reconstructing the side chains. The total runtime for creating the 202 3D models, including the time required for MODELLER, was measured. The resulting 3D models were subsequently evaluated.

*PANDORA* is a 3D modeling approach, specialized to predict the structure of pMHC complexes for both class I and II. It does so by using a database of MHC templates and a multiple sequence alignment of their conserved domains. It uses homology modeling to build the model and keeps the peptide’s anchors restrained. To evaluate the performance of PANDORA^12^ structure predictions, we made it generate pMHC-I 3D models for each of the 202 available X-ray structures. PANDORA was configured to output 20 3D models per case (the default setting) and the 3D model with the lowest molpdf score was selected. To prevent data leakage, templates containing peptides identical to those in the target 3D model were excluded from the 3D modeling process. To properly measure the process time consumption, PANDORA was executed within a multiprocessing pool configuration with 32 cores. The total processing time required for the pool to complete all 202 3D models was recorded.

*ApeGen 2.0* is a tool that generates an ensemble of peptide-MHC conformations within a chosen number of iterations. In each iteration, it first searches the Protein Data Bank (PDB) for a suitable MHC template, then anchors the atoms of the first and last two residues of the given peptide as in the template. Between these anchor positions, it samples for peptide backbone conformations that geometrically fit best using Random Coordinate Descent (RCD)^37^. On the resulting backbone conformations, OpenMM PDBFixer^30^ samples for side chains to complete the structure. Finally SMINA^38^ energy minimization is performed on both the peptide and MHC to fix steric clashes. The best resulting structure may be used as input for the next iteration.

To evaluate the structure prediction performance of APE-Gen 2.0^13^, we employed this software to 3D model the pMHC complex for each of the 202 available X-ray structures. APE-Gen 2.0 was set to output 20 3D models per case (the default setting) and the 3D model with the lowest APE-Gen 2.0 Affinity score was selected. To prevent data leakage, templates containing identical peptides to those in the target 3D model were excluded from the set. By default, an APE-Gen 2.0 process runs in a Docker container, using 8 CPU cores to pool the RCD and SMINA computations. For comparing APE-Gen 2.0’s runtime with other SOTA methods on the same software and hardware settings, we ran APE-Gen 2.0 outside a Docker container as a single process with 32 cores. The total processing time for APE-Gen 2.0 to generate all 202 3D models was recorded. These resulting 3D models were subsequently evaluated.

*MHCflurry 2.0*^7^ is a SOTA sequence-based BA predictor which consists of an ensemble of MLPs. We retrained MHCflurry 2.0 on our 7,726 BA data points by a 10-fold leave-one-cluster-out cross-validation approach for each of the 10 peptide clusters (**Suppl. Fig. 1B**). The retraining process was carried out using the allele-specific scripts, provided by MHCflurry 2.0. In each iteration, one cluster was held out as the test set, while the remaining nine clusters were used to create a training set (90%) and a network model selection set (10%). The selected network models were subsequently used to predict BA for the peptides from the remaining cluster. The predicted BA values from MHCflurry 2.0 were compared to experimental BA values (binding or non-binding) by calculating AUC. The processing time was measured.

### The training data overlap between AlphaFold2-FineTune, MHCfold and our test data

We did not retrain AlphaFold2-FineTune, because of limited time and computational resources. Instead, we utilized a pretrained network model obtained from the publicly available source (https://files.ipd.uw.edu/pub/alphafold_finetune_motmaen_pnas_2023/datasets_alphafold_finetune_v2_2023-02-20.tgz). We identified that the 202 X-ray structures in our test set represent 118 unique pMHC-I’s of which 34 were part of AlphaFold2-FineTune’s training dataset. Moreover, AlphaFold2-FineTune was fine-tuned from the AlphaFold2 model, which had been trained on PDB entries deposited before the 30th of April 2018^16^, which dates after the deposition date of 98 of those 118 unique pMHC-I’s. Therefore it is likely that these structures were included in AlphaFold2’s training dataset. That means that in total 105 of the 118 unique pMHC-I’s were used to train AlphaFold2-FineTune (see also **Fig. 4G**).

MHCfold could not be retrained for this evaluation due to the absence of a training script. Therefore, we utilized the pretrained network model available from the public source (https://github.com/dina-lab3D/MHCfold/blob/main/v7_date_5_9_2022.zip) to generate MHCfold predictions for the evaluation. For structural prediction, this model was trained on 431 unique pMHC-I’s deposited before November of 2021 with resolution higher than 3.5 Å^28^. This includes all 118 unique pMHC-I’s in our X-ray test dataset (**Fig. 4H**) and therefore it is likely that these structures were included in MHCfold’s training dataset.

For BA prediction, both the AlphaFold2-FineTune and MHCfold models were reportedly trained on the netMHCpan 4.1 training set^16,28^ which incorporates pMHC binding data from the IEDB. Since our datasets were also derived from the IEDB, it is plausible that some, if not all, of our test data was included in the training sets of these two pretrained models.

These overlaps could have inflated the results of AlphaFold2-FineTune and MHCfold.

### Evaluation of BA Prediction

We evaluated the performance of each BA predictor using the area under the receiver operating characteristic curve (AUC) as the evaluation metric. To meet the binary classification requirements of AUC, we categorized peptides as binding to their MHC when their ground truth IC₅₀ or Kd values were below 500 nanomolar (nM) and as non-binding otherwise.

### Evaluation of Structure Prediction

To calculate *Cɑ-RMSD*, all the 3D models were superposed in ProFit^39^ to the MHC G-domains of the corresponding X-ray structures. To account for structural variations, the two most N-terminal residues were omitted from the G-domain superposition due to their absence in some structures. Similarly, the C-terminal residue was excluded from the superposition analysis because its orientation varies significantly across different X-ray structures. The remaining portion of the HLA-A*02:01 sequence, encompassing the amino acids (IMGT numbering 3-179) was utilized for superposition. ProFit was used to calculate RMSD for the peptide Cα atoms to quantify the differences between the predicted 3D models and their corresponding X-ray structures. This analysis was performed for each 3D model across all methods to identify which 3D models exhibited the greatest similarity to their respective X-ray counterparts.

*Chiralities* were determined per amino acid from the positions of the N, C and Cɑ atoms around the Cα atoms using Numpy.

*Ramachandran plots* were generated by calculating the backbone ψ and φ torsion angles for each residue in the peptide using BioPython^36^.

⍵ *torsion angles* of the peptide typically approximate 180°. To verify this, the angles were calculated for each peptide bond across all 3D structures using NumPy^40^. The distributions of ω angles from the predicted 3D models and the X-ray structures were plotted to identify any differences.

*Van der Waals clashes* are considered energetically unfavorable and should be minimal. To identify overlaps between the van der Waals radii of non-bonded atom pairs in the peptide, the distances between atoms in the 3D structures were calculated using NumPy^40^. A clash was defined as occurring when two atomic centers were closer than the sum of their van der Waals radii, with exceptions made for 1) protons for which the position is usually determined by calculation, not prediction; 2) for atoms within the same residue, as those are usually either part of a ring system or connected to each other with zero, one or two atoms in between; 3) for the backbone atoms of two connected residues; 4) two connected residues where one is proline, of which the side chain is connected to the backbone; 5) two cysteines that can be disulfide bonded. Any other pair of atoms that were positioned too close to each other was considered clashing. The number and severity of clashes were compared across all 3D models.

## Data availability

The trained SwiftMHC network and related datasets (including PANDORA 3D models, BA data, torsion angles, clash counts, chiralities, RMSD and BA predictions) is accessible at with DOI 10.5281/zenodo.14968655.

## Code availability

The code for data preprocessing, training, evaluation, and prediction is available at https://github.com/X-lab-3D/swiftmhc.

## Author contributions

CB: Design and Development of the software. Running Experiments. Writing the manuscript. CG: Manuscript review and editing and code optimisation. DR: Manuscript review and editing and consultation on deep learning-related questions. David: Manuscript review, editing and testing the code. YA: Manuscript review and editing, examine possible chirality problems in AlphaFold3-generated models. LX: Design and supervise the project. Manuscript review and editing. All authors have reviewed and edited this manuscript.

## Declaration of generative AI and AI-assisted technologies in the writing process

During the preparation of this work the author(s) used ChatGPT (https://chatgpt.com) in order to improve the writing style. After using this tool, the authors reviewed and edited the content as needed and take full responsibility for the content of the publication.

## Acknowledgement

This project was supported by the Hanarth Fonds, the Kika foundation (grant number 454) and NWO-XS (OCENW.XS23.2.130). The computation was partly supported by the NVIDIA Academic Grant Program. We thank SurfSara for their generous GPU and CPU computing resources (grant numbers EINF2380, EINF-10427 and EINF-11930). We also thank Dario Marzella for providing the dataset and helpful discussions.

