## supplementary for "SwiftMHC: A High-Speed Attention Network for MHC-Bound Peptide Identification and 3D Modeling"

### 1. Data

#### 1.1 Data Composition

To examine the distribution of peptide lengths in pMHC class I (pMHC-I) complexes, we downloaded 3,064,594 MHC-I BA data entries from the IEDB[4]. These entries encompass complexes of various peptide lengths and MHC-I alleles, beyond just 9-mer peptides and the HLA-A*02:01 allele. The histogram plot indicates that 9-mer peptides occur most frequently in this database (**Suppl. Fig. 1A**).

SwiftMHC is trained, validated and tested on a subset of these 3,064,594 entries. This subset consists of 7,726 HLA-A*02:01 9-mer BA data from the IEDB[4]. Because SwiftMHC requires structural data to train on, these entries were represented by 7,726 PANDORA models. Additionally SwiftMHC was validated on 202 X-ray structures. To avoid data leakage, this data was combined and Gibbs clustered[7] based on peptide sequence similarities into 10 clusters of roughly the same size. The distribution of clustered data is shown in **Suppl. Fig. 1B**.

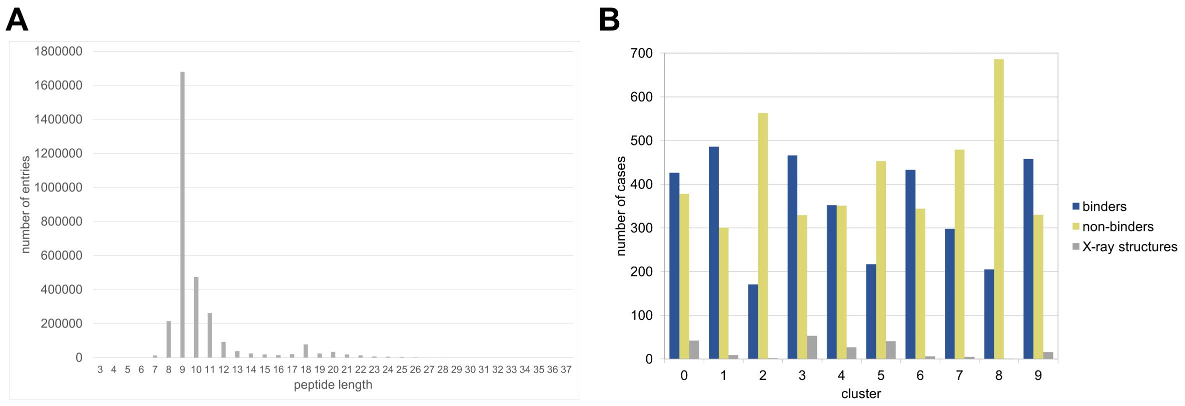

**Supplementary Figure 1: Data distributions. A.** Distribution of pMHC-I peptide lengths in IEDB[4] (as of 26/11/2024). In total, 3,064,594 entries were counted. These represent all occurring lengths and MHC-I alleles, not just 9-mers and HLA-A*02:01. **B.** Distribution of binders, non-binders and X-ray structures across clusters. IEDB binders: $K_d$ or $IC_{50} < 500$ nM; non-binders: $K_d$ or $IC_{50} > 500$ nM.

#### 1.2 Data Preprocessing

Before SwiftMHC can be trained from the input data (**Subsection 1.1**) it is preprocessed. During preprocessing, several variables are precalculated and stored in HDF5 (https://www.hdfgroup.org/solutions/hdf5/) format with LZMA compression (http://www.7zip.org/7z.html) for every pMHC complex: *ground truth frames, ground truth torsion angles, a MHC proximity matrix, amino acid sequences, sequence masks* and *ground truth BA*. This section describes each of these in detail. For ground truth structural data, either X-ray 3D structures from the RCSB PDB[5] or

PANDORA[3] 3D models were used. Ground truth BA data were extracted from the IEDB[4].

##### 1.2.1 Ground Truth Frames for Backbone and Side Chain Conformations

The *geometry* of backbone and side chains of both the MHC and peptide are represented by local frames (or local coordinate frames). Local frames are attached to objects (amino acids here) to describe their orientation and position relative to a global frame (**Suppl. Fig. 2A**). This global frame represents the xyz-coordinate system of the pMHC structure. The local frames allow us to describe the movement of residues or side-chains. Each local frame $T_i$ holds a translation vector $\vec{t}_i$ and a unit quaternion $q_i$ that describes a rotation: so $T_i = (q_i, \vec{t}_i)$.

Applying a transformation to global space means that the point vector is first rotated by the quaternion and then translated by the corresponding translation vector. In this document, such a frame transformation is indicated by the ∘ operator.

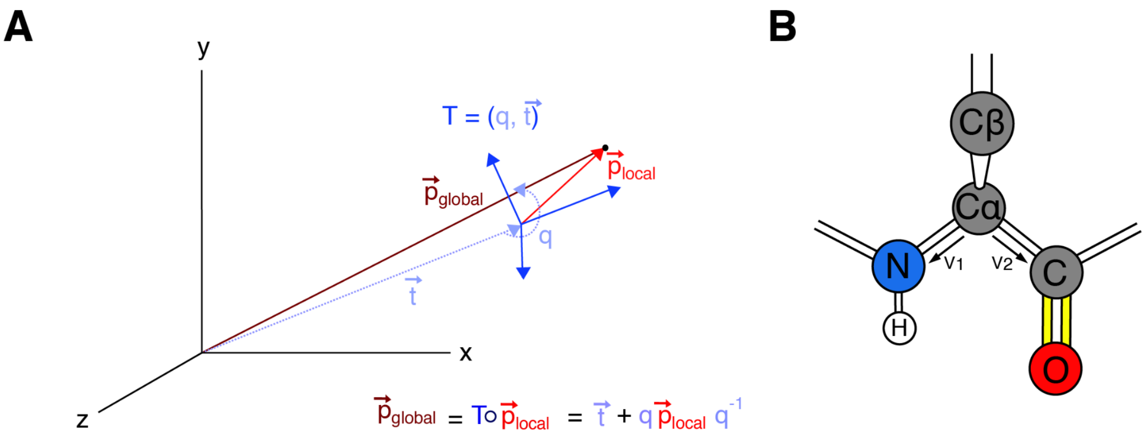

**Supplementary Figure 2: Frames.** Local frames are coordinate systems attached to each amino acid to define their position and orientation relative to the global reference coordinate system. In **A.** a point $\vec{p}$ in the local frame space (blue axes) can be converted to the global space (black axes) by transforming it by the frame *T*, consisting of a rotation quaternion *q* and a translation vector $\vec{t}$. Such a transformation is indicated by the ∘ operator. **B.** displays how a backbone local frame for one residue is defined from the atomic positions of backbone N, Cα and C. The bond between atoms N and Cα ($\vec{v}_1$) determines the direction of the x-axis and the three atoms together lie in the xy-plane ($\vec{v}_1$ and $\vec{v}_2$), from which the y-axis is derived. The z-axis is the normalized cross product between the x and y axis and is perpendicular to this xy-plane, $\vec{v}_1$ and $\vec{v}_2$.

In order to predict structure, SwiftMHC predicts frames, but it also takes frames as ground truth during training. *Ground truth frames* are derived from 3D protein structures and include translations relative to a defined origin. To ensure consistency across all processed structures, the origin was set to the center of the MHC groove, and all structures were superposed accordingly. Additionally, the superposed structures were aligned to share the same orientation.

During preprocessing, the 7 ground truth frames and torsion angles are calculated from the atomic positions of each residue. These frames are always defined from three atomic positions, where $\vec{t}_i$ is the position of one of the atoms and $q_i$ is calculated to represent the orientation of the other two atoms around this central atom (**Suppl. Fig 2B**). For the backbone truth frames $\{T_i^{true}\}$, we use the N,Cα,C atoms. The seven side chain ground truth frames $\{T_i^{true,\, sidechain}\}$ are based on rotatable covalent bonds, that define the conformation of an amino acid by at most seven torsion angles $\varphi$, $\psi$, $\omega$, $\chi_1$, $\chi_2$, $\chi_3$, $\chi_4$. Specifically, we use the two atoms of each torsion bond and the next atom after that. Because for some torsion angles in some amino acids there is a 180° symmetry in the side chain, we also compute a series of alternative torsion angles and frames to be stored in separate arrays: $\{T_i^{alt\ truth}\}$ and $\{T_i^{alt\ truth,\, sidechain}\}$. For smaller amino acids with less than 7 torsion angles, the nonexistent frames and angles are masked out to be ignored by the algorithm.

##### 1.2.2 The MHC Proximity Matrix

For every residue pair in the MHC, the distance between the closest two heavy atoms is determined. These distance values $d_{j\hat{j}}$ are converted to proximity values $z_{j\hat{j}}$ by the formula: $z_{j\hat{j}} = \frac{1}{1 + d_{j\hat{j}}}$. These values are combined into an $r_j \times r_j \times 1$ proximity matrix, where $r_j$ is the number of residues in the MHC G-domain. This matrix $z_{j\hat{j}}$ is input to the MHC Self IPA module (**Subsection 2.2**).

##### 1.2.3 Amino Acid Sequences

Peptide and MHC sequences are represented as one-hot encodings, where the corresponding number of the amino acid is set to 1 and the other numbers are set to 0. Each of the 20 amino acid types is represented by a one-hot encoded $c_s \times 1$ vector with $c_s = 32$ and therefore a sequence of $r$ residues is represented by a $r \times c_s$ array of vectors. For the dimension $c_s$, the encoding scheme uses a number larger than 20 to allow flexibility for representing non-canonical amino acids. This approach ensures compatibility with potential future extensions involving non-standard amino acid residues, such as post-translationally modified variants.

Additionally each amino acid in a sequence is associated with an atomic mask and a list of atom names. Atomic masks are essential for structural loss functions, as they specify which xyz positions should be included in the calculations. Atom names are necessary for converting arrays of xyz positions into actual atoms when generating PDB-formatted 3D structures.

##### 1.2.4 Sequence Masks

For computational efficiency, SwiftMHC works with masked MHC structures, meaning that a selection is made for residues to be included in the attention modules. For MHC Self IPA calculation (**Subsection 2.2**), only residues within the G-domain (IMGT[9] numbers 2-180) are used. For Cross IPA between peptide and MHC (**Subsection 2.4.1**), only MHC residues within 10 Å of the peptide (as observed in any X-ray structure) are included (**Suppl. Fig. 3**).

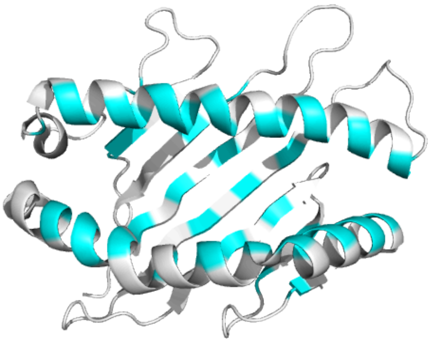

**Supplementary Figure 3: HLA-A*02:01 residues that participate in Cross IPA (cyan).** PDB entry 3MRD is used here and the peptide is not shown for easy visualisation.

##### 1.2.5 Ground Truth BA

From the IEDB[4] experimental BA data, both $IC_{50}$ and $K_d$ values are converted to $1 - log_{50000}(IC_{50})$ or $1 - log_{50000}(K_d)$ respectively as was done in [10]. The resulting values serve as the ground truth in the BA loss function in regression-based BA training and validation.

The class label (binding or non-binding) is also derived from this BA value: pMHC complexes are considered binding if their $IC_{50}$ or $K_d$ is below 500 nM. This class label is used for calculating the receiver operating characteristic area under the curve (AUC).

#### 1.3 Data Representation

SwiftMHC's main module predicts peptide 3D structures and BA values (**Subsection 2.1**). The module processes preprocessed input data, including the MHC and peptide *sequences*, the *local backbone frames* of the MHC, and a *proximity matrix* specific to the MHC.

Initially the peptide and MHC are represented by two *sequences* of one-hot encoded amino acids (**Subsection 1.2.3**). We denote the peptide sequence as $\{s_i\}$ and the MHC sequence as $\{s_j\}$. These vectors serve as input to the Self Attention modules (**Algorithms 2** and **3**) that will output updated versions of them. We denote the array of *local backbone frames* as $\{T_i\}$ for the peptide and $\{T_j\}$ for the MHC. The MHC frames $\{T_j\}$ are set to be equal to the ground truth (**Subsection 1.2.1**) and they are kept fixed during inference. The backbone frames for the peptide $\{T_i\}$ are initially all placed at the center of the MHC groove with uniform orientation, but they are iteratively refined during inference to approach their ground truth. Specifically to the MHC an $r_j \times r_j \times 1$ *proximity matrix* $\{z_{j\hat{j}}\}$ (**Subsection 1.2.2**) is used. This matrix serves as input to the MHC Self Invariant Point Attention Module (**Subsection 2.2**).

The output of the Main module consists of a numerical BA value and a peptide structure. This peptide structure is represented by the xyz positions of the peptide atoms themselves. These are represented by a $r_i \times n \times 3$ array of vectors $\{\vec{x}_i^{all}\}$, where $r_i$ represents the number of residues in the peptide sequence and $n$ represents the maximum number of atoms per residue ($n = 14$). For small amino acids like alanine, not all of these vectors are used. Atomic masks are used in the loss function to make sure that the positions of nonexistent atoms do not count.

#### 2 SwiftMHC Algorithm Design

##### 2.1 The Main module

The neural network of SwiftMHC consists of a Main module (**Algorithm 1**). This module takes as input a major histocompatibility complex (MHC) structure and a peptide sequence. Its outputs are the *BA* value and an all-atom pMHC 3D model.

**Algorithm 1** MainModule

**procedure** MainModule ( $\{s_i\}, \{s_j\}, \{z_{j\hat{j}}\}, N_{blocks} = 2,$
$\{T_j\}, \{T_i^{true,all}\}, \{T_i^{alt\ truth,all}\},$
$\{\vec{x}_j^{all}\}, \{\vec{x}_i^{true,all}\}, \{\vec{x}_i^{alt\ truth,all}\},$
$\{\vec{\alpha}_i^{true,all}\}, \{\vec{\alpha}_i^{alt\ truth,all}\}$,BA$^{true}$ ):

`# This layer applies weighted normalization over all pairs`
`# of MHC residues.  It is likely to be removed in future updates.`
$\{z_{j\hat{j}}\} \leftarrow \text{LayerNorm}(\{z_{j\hat{j}}\})$

`# MHC self attention (module 1).`
`# SelfIPA and LayerNorm weights are shared across blocks.`
`#` $j$ `and` $\hat{j}$ `correspond to MHC residues.`
**for all** $l \in [1, ..., N_{blocks}]$ **do**
    `# For SelfIPA see Algorithm 2.`
    $\{s_j\} += \text{SelfIPA}(\{s_j\}, \{z_{j\hat{j}}\})$
    $s_j \leftarrow \text{LayerNorm}(s_j)$
**end for**

`# Peptide self attention (module 2).`
`# Each block has its own set of PeptideSelfAttention weights and biases.`
`# LayerNorm weights are shared across blocks.`
`#` $i$ `corresponds to a peptide residue.`
**for all** $l \in [1, ..., N_{blocks}]$ **do**
    `# For PeptideSelfAttention see Algorithm 3.`
    $\{s_i\} += \text{PeptideSelfAttention}(\{s_i\})$
    $s_i \leftarrow \text{LayerNorm}(s_i)$
**end for**

`# Perform Cross Attention (module 3) and predict structure.`
`# For CrossAttentionStructureModule see Algorithm 4.`
$\{\vec{x}_i^{all}\}, \{T_i^{all}\}, \{\vec{\alpha}_i^{all}\}, \{s_i\} =$
    CrossAttentionStructureModule ( $\{s_i\}, \{s_j\}, \{T_j\}$ )

`# Predict BA (module 4).`
`# BA` $\in \mathbb{R}$
`# For PredictBindingAffininty see Algorithm 8.`
BA =PredictBindingAffinity( $\{s_i\}$)

38:

39:     `# Loss computation is optional.`

40:     `# For CalculateLoss, see Algorithm 9`

41:     $\mathcal{L}_{\text{tot}} = \text{CalculateLoss}(\text{BA}, \text{BA}^{true},$

42:         $\{\vec{x}_i^{all}\}, \{\vec{x}_j^{all}\}, \{\vec{x}_i^{true,all}\}, \{\vec{x}_i^{alt\ truth,all}\},$

43:         $\{T_i^{all}\}, \{T_i^{true,all}\}, \{T_i^{alt\ truth,all}\},$

44:         $\{\vec{\alpha}_i^{all}\}, \{\vec{\alpha}_i^{true,all}\}, \{\vec{\alpha}_i^{alt\ truth,all}\}\ )$

45:

46:     **return** BA, $\{\vec{x}_i^{all}\}$, $\mathcal{L}_{tot}$

47: **end procedure**

---

##### 2.2 Module 1: MHC Self Invariant Point Attention

Within the Main module, the MHC Self Invariant Point Attention (Self IPA) module operates on the MHC structural data (**Suppl. Fig. 4, Algorithm 2**). The purpose of this module is to encode each MHC residue-based on its amino acid type and its structurally neighboring amino acids. This module draws inspiration from **AlphaFold2**[1] **Suppl. Algorithm 22 - Invariant Point Attention,** which enables amino acid residues in a sequence to update each other based on their features and structural geometry. The code for this module is a modified version of the Openfold[11] IPA implementation. It was designed to represent and process interactions between amino acids. The inputs of the module are the one-hot encoded MHC amino acid sequence $\{s_j\}$ and the proximity matrix $\{z_{j\hat{j}}\}$. For computational efficiency purposes, the MHC structure is masked so that only residues of the G-domain are included in the computation. The computed attention weights $a_{j\hat{j}}^h$ are used for updating the MHC residue features. The output of this submodule is a sequence of updated feature vectors $\{s_j\}$ that is passed on to the next block. This Self IPA submodule is called in two iterative blocks so that the output of the previous block is the input to the next. A skip connection was added in each block (**Algorithm 1, line 15**) to overcome potential degradation problems. The learned Self IPA weights are shared across blocks. The output of the final block is one of the input variables in the Cross Attention Structure module (**Algorithm 1, line 31-32**).

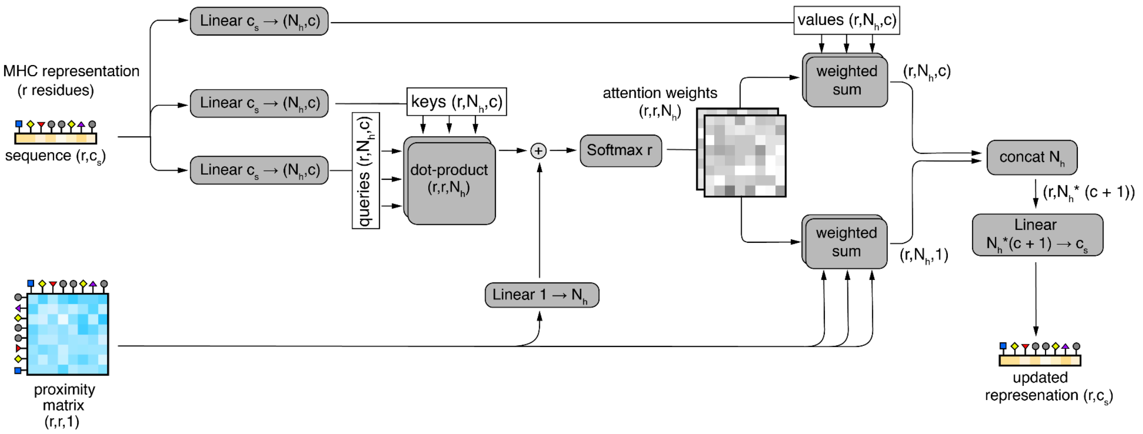

**Supplementary Figure 4**: **Flow chart of the MHC Self IPA module.** Dimensions: r: MHC length (r ≤ 200), $c_s$: amino acid input channels ($c_s$ = 32), c: hidden channels (c = 16), $N_h$: number of heads ($N_h$ = 2). We use ⊕ for element-wise addition.

---

**Algorithm 2** Self Invariant Point Attention

---

**procedure** SelfIPA ( $\{s_j\}, \{z_{j\hat{j}}\}, c = 16, N_h = 2$ ) :

    # Here, $j$ and $\hat{j}$ correspond to residues in the MHC.
    # $q_j^h, k_j^h, v_j^h \in \mathbb{R}^c, h \in \{1, ..., N_h\}$
    $q_j^h, k_j^h, v_j^h = \text{LinearNoBias}(s_j)$
    $b_{j\hat{j}}^h = \text{LinearNoBias}(z_{j\hat{j}})$

    # Calculate attention weights $\{a_{j\hat{j}}^h\}$.
    $w_L = \sqrt{\frac{1}{2}}$
    $a_{j\hat{j}}^h = \text{softmax}_{\hat{j}}(w_L(\frac{1}{\sqrt{c}} q_j^{h\top} k_{\hat{j}}^h + b_{j\hat{j}}^h))$

    # Apply weighted attention.
    $o_j^h = \sum_{\hat{j}} a_{j\hat{j}}^h v_{\hat{j}}^h$
    $\tilde{o}_j^h = \sum_{\hat{j}} a_{j\hat{j}}^h z_{j\hat{j}}$

    $s_j \leftarrow \text{Linear}(\text{concat}_h(o_j^h, \tilde{o}_j^h))$
    **return** $s_j$
**end procedure**

---

#### 2.3 Module 2: Peptide Self Attention

Within the Main module, the Peptide Self Attention module operates on peptide sequential data (**Algorithm 3** and **Suppl. Fig. 5**). The peptide structure is unknown at this point and this module updates the feature representation of each peptide residue based on its own amino acid type and its sequential neighbors. We intend to design an algorithm that can eventually model peptides with variable lengths other than just 9. For this reason we employed relative position encoding (**Algorithm 3,**

**lines 2-7, Suppl. Fig. 5B**). Using the outer differences between residue positions $f_i^{residue\_index}$, we construct a relative positions matrix, where each element $d_{i\hat{\imath}}$ represents the distance between two residues $i$ and $\hat{\imath}$, measured as negative for N-terminal distances and positive for C-terminal distances (**Algorithm 3, line 6**). These differences are calculated and one-hot encoded into $c_z = 33$ different bins ($v_{bins} = [-16, -15, ...0...15, 16]$), forming the relative position encodings $\{z_{i\hat{\imath}}\}$ in **Algorithm 3.** Those relative position encodings are used in the peptide Self Attention computation.

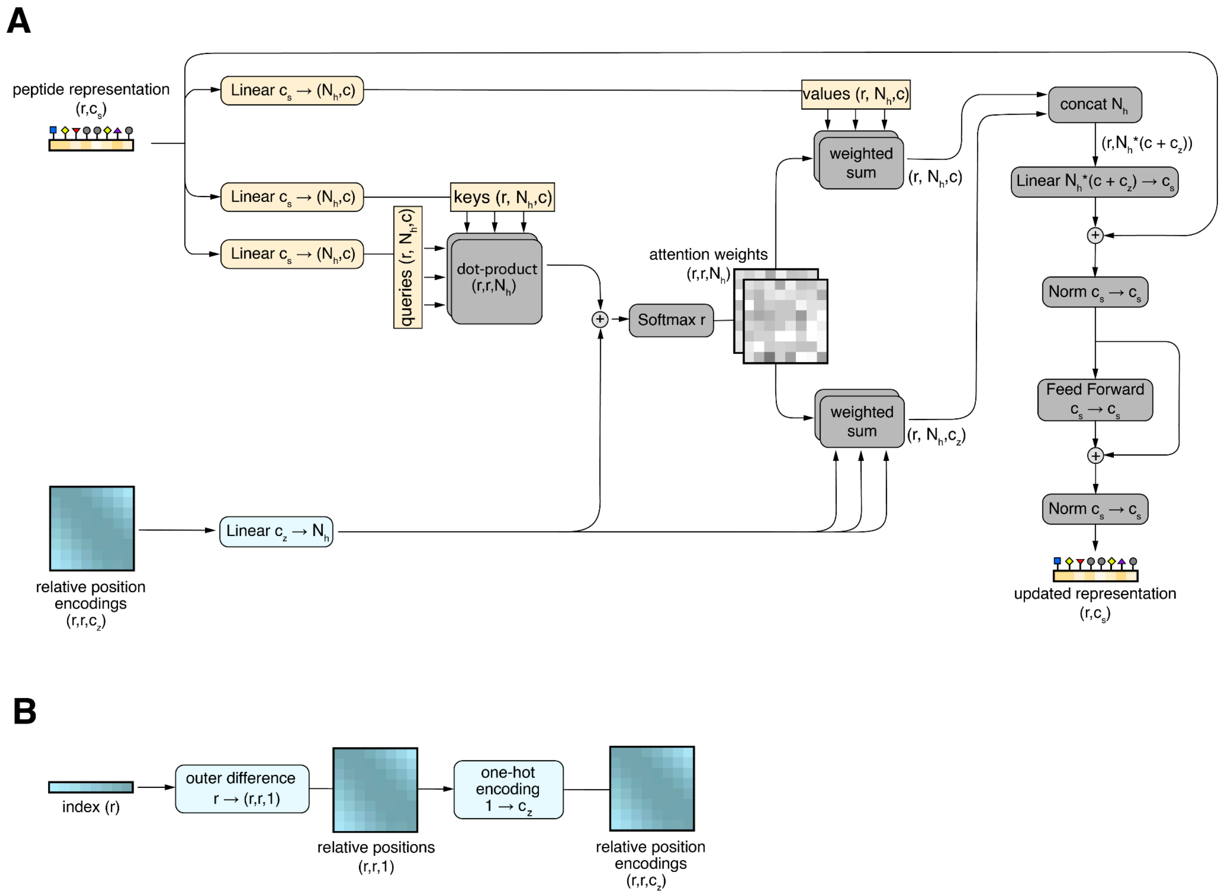

**Supplementary Figure 5**: **Flow chart of the Peptide Self Attention module.** Dimensions: r: peptide length ( r ≤ 16), $c_s$: amino acid input channels ($c_s$ = 32), $c_z$: relative position bins ($c_z$=33), c: hidden channels (c=16), $N_h$: heads ($N_h$=2). We use ⊕ for element-wise addition. **A.** Full module. **B.** Relative position encoding.

The peptide Self Attention is computed similarly to regular multi-headed attention[2]. From the the one-hot encoded input sequence $\{s_i\}$, the queries($q_i^h$), keys($k_i^h$) and values($v_i^h$). The queries and keys are dot multiplied, normalized and then used as the first term for the multi-headed attention weights. The key difference from regular multi-headed attention is that the formula for attention weights includes a second term, additionally to the normalized dot product (**Algorithm 3, line 17**).

This second attention term $\{b^h_{i\hat{i}}\}$ is derived from the relative position encodings $\{z_{i\hat{i}}\}$. A trainable linear weight matrix converts them into $\{b^h_{i\hat{i}}\}$. Softmax is used to convert the sum of the two terms into actual attention weights $\{a^h_{i\hat{i}}\}$, summing up to 1.0 over each peptide residue. (**Algorithm 3, line 17**).

The weights in $\{a^h_{i\hat{i}}\}$ are multiplied by the values $\{v^h_i\}$ and by one-hot encoded relative position encodings $\{z_{i\hat{i}}\}$. The results of those multiplications are concatenated, embedded and then inserted into a feed forward module (**Algorithm 3, lines 24,25**). This feed forward module was placed here from inspiration of the PyTorch transformer encoder[3].

The Peptide Self Attention module is called in two iterative blocks (**Algorithm 1, lines 23-27**). The result of one block altogether is an updated version of the sequence embedding: $\{s_i\}$ (**Algorithm 3, line 27**). The output of the previous block is the input to the next block. A skip connection was added (**Algorithm 1, line 26**) in each block to overcome potential degradation problems. The output of the final block is used as input to the Cross Attention Structure module (**Algorithm 1, line 31-32**). The learned Peptide Self Attention module's weights are not shared across blocks. Instead, each block maintains its own distinct set of weights.

**Algorithm 3** Peptide Self Attention

1: **procedure** PeptideSelfAttention( $\{s_i\}, c = 16, c_z = 33, N_h = 2$ ):
2: `# Relative Position Encoding:`
3: `#` $i$ `and` $\hat{i}$ `correspond to peptide residues.`
4: $f_i^{residue\_index} = i$
5: $v_{bins} = [-16, -15, ..., 16]$
6: $d_{i\hat{i}} = f_i^{residue\_index} - f_{\hat{i}}^{residue\_index}$
7: $z_{i\hat{i}} = \mathrm{one\_hot}(\mathrm{argmin}(|d_{i\hat{i}} - v_{bins}|))$
8:
9: `#` $z_{i\hat{i}} \in \mathbb{R}^{c_z}, b^h_{i\hat{i}} \in \mathbb{R}, h \in \{1, ..., N_h\}$
10: $b^h_{i\hat{i}} = \mathrm{LinearNoBias}(z_{i\hat{i}})$
11:
12: `#` $q^h_i, k^h_i, v^h_i \in \mathbb{R}^{c_h}$
13: $q^h_i, k^h_i, v^h_i = \mathrm{LinearNoBias}(s_i)$
14:
15: `# Calculate attention weights` $\{a^h_{i\hat{i}}\}$.
16: $w_L = \sqrt{\frac{1}{2}}$
17: $a^h_{ij} = \mathrm{softmax}_{\hat{i}}(w_L(\frac{1}{\sqrt{c}} q^h_i k^h_{\hat{i}} + b^h_{i\hat{i}}))$

`# Apply weighted attention.`

$\tilde{o}_i^h = \sum_{\hat{i}} a_{i\hat{i}}^h z_{i\hat{i}}$

$o_i^h = \sum_{\hat{i}} a_{i\hat{i}}^h v_{\hat{i}}^h$

`# Skip connection before feed-forward module.`

$s_i += \mathrm{Linear}(\mathrm{concat}_h(\tilde{o}_i^h, o_i^h))$

$s_i \leftarrow \mathrm{LayerNorm}(\mathrm{Dropout}_{0.1}(s_i))$

`# Feed-forward:`

$s_i += \mathrm{Linear}(\mathrm{relu}(\mathrm{Linear}(s_i)))$

$s_i \leftarrow \mathrm{LayerNorm}(\mathrm{Dropout}_{0.1}(s_i))$

**return** $\{s_i\}$

**end procedure**

##### 2.4 Module 3: Cross Attention Structure module

The Cross Attention Structure module (**Algorithm 4**) is called from inside the Main module, after executing the two Self Attention modules. The Cross Attention Structure module was named after the Structure module in AlphaFold2[1], which inspired its design. It is called *Cross Attention* because it applies a form of multi-headed attention between the MHC and the peptide. The purpose of this module is to predict a structure for the peptide, given peptide sequence features $\{s_i\}$ and an MHC protein, represented by its sequence features $\{s_j\}$ and backbone structure frames $\{T_j\}$ (**Subsection 1.2.1**). This module also returns an updated peptide representation $\{s_i\}$ that contains sequence and structural information of the predicted pMHC structure. This updated representation is subsequently passed to the BA module for BA predictions. Two iterative blocks are used (**Algorithm 4, lines 15-53**) to consider a longer range of residue attentions. Each block contains multiple skip connections to overcome potential gradient degradation problems.

In each block, the module updates both the peptide's feature representation $\{s_i\}$ and its backbone frames $\{T_i\}$. Cross Invariant Point Attention (*Cross IPA*) is applied first (**Subsection 2.4.1**), using MHC information to update $\{s_i\}$. Cross IPA is followed by a transitional module, a dropout operation and a normalization layer (**Algorithm 4, lines 28,29**), the purpose of which are to make $s_i$-encoded information more expressive. Next, the $\{s_i\}$ variable is used as input for updating $\{T_i\}$ (**Subsection 2.4.2)**. From $\{T_i\}$, the positions of the Cα atom and positions of its directly connected atoms N, C, Cβ will be calculated to be saved in the atomic position array $\{\vec{x}_i^{all}\}$. The next step is to predict the side chain torsion angles $\chi_1$, $\chi_2$, $\chi_3$, $\chi_4$ from the updated $\{s_i\}$ as input. The side chain frames $\{T_i^{sidechain}\}$ are calculated from $\{T_i\}$ and the predicted side chain torsion angles. From $\{T_i^{sidechain}\}$ and the corresponding rigid groups (**AlphaFold2**[1] **Suppl. Table 2**), all other atom positions are calculated. Finally, the ω torsion angles are calculated from the resulting $C\alpha_i$-$C_i$-$N_{i+1}$-$C\alpha_{i+1}$ positions (**Algorithm 7**). The loss function will compare those ω angles with the ground truth later (**Suppl. Fig.7**). The resulting loss values will be used for feedback in training the network. This additional computation step was added to boost accuracy of the predicted ω angle output.

Once a Cross Attention Structure module block is completed, the resulting $\{s_i\}$ and $\{T_i\}$ are used as input for the next block. The output of the Cross Attention Structure module are the updated peptide features $\{s_i\}$, the local frames $\{T_i^{all}\}$, atomic positions $\{\vec{x}_i^{all}\}$ and torsion angles $\{\vec{\alpha}_i^{all}\}$ from the final block. The output peptide features $\{s_i\}$ will be used as input for the SwiftMHC BA Predictor (**Subsection 2.5**).

**Algorithm 4** Cross Attention Structure module

**procedure** CrossAttentionStructureModule $(\{s_i^{initial}\}, \{s_j^{initial}\},$
$\{T_j\}, N_{blocks} = 2)$:
    # $i$ corresponds to a peptide residue.
    # $j$ corresponds to a MHC residue.
    $s_i^{initial} \leftarrow \mathrm{LayerNorm}(s_i^{initial})$
    $s_j^{initial} \leftarrow \mathrm{LayerNorm}(s_j^{initial})$

    # $T_i$ is initialized as a series of identity frames, this works well
    # only when the preprocessing superimposed
    # all input structures in the same orientation.
    $q_I = (1,0,0,0)$ # quaternion definition:  q = (w,x,y,z)
    $\vec{t}_I = (0,0,0)$
    $T_i = (q_I, \vec{t}_I)$

**for all** $l \in [1, ..., N_{blocks}]$ **do**

`# Cross Invariant Point Attention (CrossIPA)`
`# between peptide and MHC:`

`# The weights of the cross IPA are shared across blocks.`
`# For CrossIPA see Algorithm 5.`
$\{s_i\} += \mathrm{CrossIPA}(\{s_i\}, \{s_j\}, \{T_i\}, \{T_j\})$
$s_i \leftarrow \mathrm{LayerNorm}(\mathrm{Dropout}_{0.1}(s_i))$

`# Structure module transition:`

`# The weights of the Linears and LayerNorm`
`# are shared across blocks.`
$s_i \leftarrow s_i + \mathrm{Linear}(\mathrm{relu}(\mathrm{Linear}(\mathrm{relu}(\mathrm{Linear}(s_i)))))$
$s_i \leftarrow \mathrm{LayerNorm}(\mathrm{Dropout}_{0.1}(s_i))$

`# BackboneUpdate is similar to AlphaFold2`[1] `Suppl. Algorithm 23.`
`# It updates the backbone frames by transformation as`
`# described in AlphaFold2`[1] `Suppl. Subsection 1.1`

`# All weights of BackboneUpdate are shared across blocks.`
`# For BackboneUpdate, see Algorithm 6.`
$T_i \leftarrow T_i \circ \mathrm{BackboneUpdate}(s_i)$

`# Predict side chain and backbone angles` $\phi, \psi, \omega, \chi_1, \chi_2, \chi_3, \chi_4$
`# as sin,cos vectors.`
`# All weights of side chain angle prediction`
`# are shared across blocks.`
$a_i = \mathrm{Linear}(\mathrm{relu}(s_i)) + \mathrm{Linear}(\mathrm{relu}(s_i^{initial}))$
$a_i \leftarrow a_i + \mathrm{Linear}(\mathrm{relu}(\mathrm{Linear}(\mathrm{relu}(a_i))))$
$a_i \leftarrow a_i + \mathrm{Linear}(\mathrm{relu}(\mathrm{Linear}(\mathrm{relu}(a_i))))$
$\vec{\alpha}_i^{all} = \mathrm{normalize\_vec}(\mathrm{Linear}(\mathrm{relu}(a_i)))$

47: # Rotation gradients are disabled between blocks,
48: # to stabilize training.
49: **if** $l < N_{block}$ **then**
50: $q_i, \vec{t_i} \leftarrow T_i$
51: $T_i \leftarrow (\text{stopgrad}(q_i), \vec{t_i})$
52: **end if**
53: **end for**
54:
55: # ComputeAllAtomCoordinates is from AlphaFold2[1] Suppl. Algorithm 24.
56: $T_i^{sidechain}, \vec{x}_i^{all} = \text{ComputeAllAtomCoordinates}(T_i, \vec{\alpha}_i^{all})$
57: $T_i^{all} = \text{concat}(T_i, T_i^{sidechain})$
58:
59: # Update the peptide $\omega$ angles from the computed backbone
60: # coordinates.
61: # For CalculateTorsion, see Algorithm 7.
62: $(\vec{\omega}_i, \vec{\phi}_i, \vec{\psi}_i, \vec{\chi_1}_i, ...) = \vec{\alpha}_i^{all}$
63: $\{\vec{x}_i^N, \vec{x}_i^{C\alpha}, \vec{x}_i^C, ...\} = \vec{x}_i^{all}$
64: $\{\vec{x}_{i+1}^N, \vec{x}_{i+1}^{C\alpha}, \vec{x}_{i+1}^C, ...\} = \vec{x}_{i+1}^{all}$
65: $\vec{\omega}_i = \text{CalculateTorsion}(\vec{x}_i^{C\alpha}, \vec{x}_i^C, \vec{x}_{i+1}^N, \vec{x}_{i+1}^{C\alpha})$
66: $\vec{\alpha}_i^{all} = (\vec{\omega}_i, \vec{\phi}_i, \vec{\psi}_i, \vec{\chi_1}_i, ...)$
67:
68: **return** $\{\vec{x}_i^{all}\}, \{T_i^{all}\}, \{\vec{\alpha}_i^{all}\}, \{s_i\}$
69: **end procedure**

##### 2.4.1 Cross Invariant Point Attention

The Cross Invariant Point Attention (Cross IPA) module (**Algorithm 5** and **Suppl. Fig. 6**) is part of the Cross Attention Structure module (**Algorithm 4, line 21**). SwiftMHC uses the Cross IPA module to update the peptide representation $\{s_i\}$ using the predicted $\{T_i\}$ and information from the MHC groove, namely $\{s_j\}$ and the invariable $\{T_j\}$. Peptide residue representations serve as queries, and MHC representations as keys and values in the attention mechanism. The updated peptide representation $\{s_i\}$ will be used to update the coordinates of peptide residues $\{T_i\}$ and to be passed to the BA module to predict BA.

Cross IPA is a multi-headed Cross Attention module that takes into account both the sequence features and geometry. Thus the attention weights have two components: sequence-level $q_i^{h\top} k_j^h$ and geometric level $||T_i \circ \vec{q}_i^{hp} - T_j \circ \vec{k}_j^{hp}||$ (**Algorithm 5**, **Line**

**22**). The first term $q_i^{h\top} k_j^h$ calculates the attention weights between each peptide residue and each MHC residue in terms of amino acid similarities. To calculate the geometric attention, 4 virtual points are created for each peptide residue and 4 virtual points are created for each MHC residue (**Algorithm 5 Line 11-13**). The distance $||T_i \circ \vec{q}_i^{hp} - T_j \circ \vec{k}_j^{hp}||$ is calculated between a virtual point of a peptide residue and a virtual point of a MHC residue. The virtual points are initially placed in the backbone local frame space (see **Subsection 1.2.1** for explanation) and the peptide $\{T_i\}$ and MHC $\{T_j\}$ are used to transform those virtual points to global space. The global space coordinates of those virtual points are used to calculate the squared distances that are summed up to form the second term in the attention weights calculation (**Algorithm 5, line 22**).

The Cross Invariant Point Attention module does not operate on the entire MHC G-domain. For speed purposes, only a masked selection of MHC residues is used. We selected all residues that are within a 10 Å radius from the peptide in any of the known pMHC-I structures (**Suppl. Figure 3).**

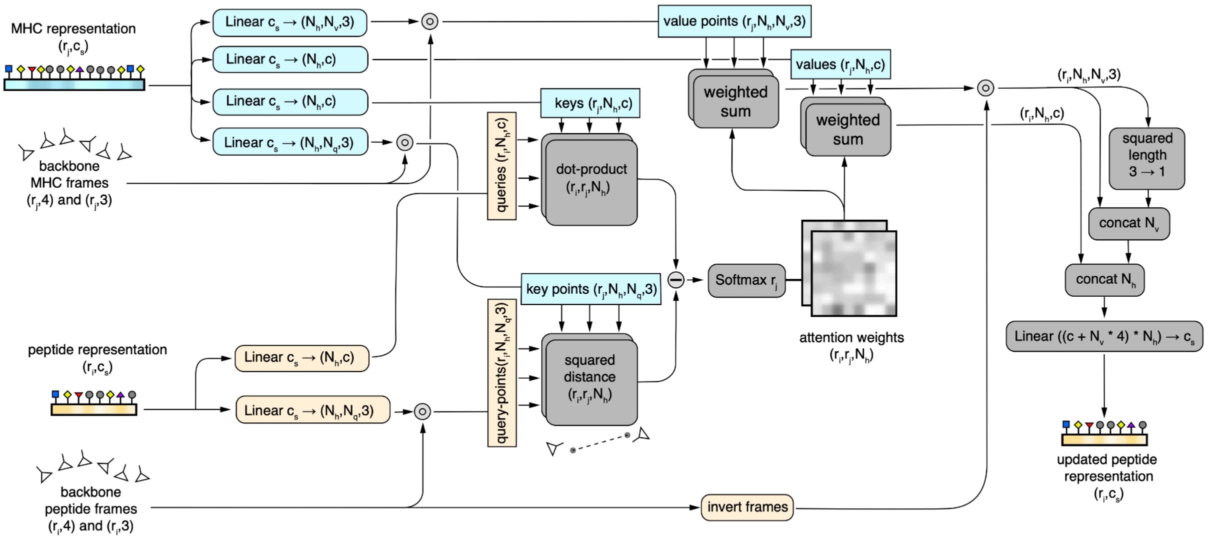

**Supplementary Figure 6: Flow chart of the Cross IPA module.** Dimensions: $r_i$: peptide length ($r_i \leq 16$), $r_j$: MHC length ($r_j \leq 200$), $c_s$: amino acid input channels ($c_s = 32$), $N_q$ : number of generated query and key points($N_q = 4$), $N_v$: number of generated value points ($N_v = 8$) , c: hidden channels (c = 16), $N_h$: heads ($N_h = 2$). We use ⊚ for transforming vectors by frames and ⊖ for element-wise subtraction. The two triangles represent two local frames: one frame represents a peptide residue and transforms query points; the other frame represents a MHC residue and transforms key points. A dashed line indicates distance measured between those two invariant points predicted within the frame-local space.

**Algorithm 5** Cross Invariant Point Attention

**procedure** CrossIPA ( $\{s_i\}, \{s_j\}, \{T_i\}, \{T_j\}, \{T_i\}$
$c = 16, N_h = 2, N_q = 4, N_v = 8$ ):

    # $i$ corresponds to a peptide residue.
    # $j$ corresponds to a MHC residue.

    # $q_i^h, k_j^h, v_j^h \in \mathbb{R}^c, h \in \{1, ..., N_h\}$
    $q_i^h = LinearNoBias(s_i)$
    $k_j^h = LinearNoBias(s_j)$
    $v_j^h = LinearNoBias(s_j)$

    # $\vec{q}_i^{hp}, \vec{k}_j^{hp} \in \mathbb{R}^3, p \in \{1, ..., N_q\}$
    $\vec{q}_i^{hp} = LinearNoBias(s_i)$
    $\vec{k}_j^{hp} = LinearNoBias(s_j)$

    # $\vec{v}_j^{hp} \in \mathbb{R}^3, p \in \{1, ..., N_v\}$
    $\vec{v}_j^{hp} = LinearNoBias(s_j)$

    # Calculate attention weights $\{a_{ij}^h\}$ .
    # The head weights $\{\gamma^h\}$ are trainable.
    $w_C = \sqrt{\frac{2}{9N_q}}$
    $w_L = \sqrt{\frac{1}{2}}$
    $a_{ij}^h = \mathrm{softmax}_j(w_L(\frac{1}{\sqrt{c}} q_i^{h\top} k_j^h - \frac{\gamma^h w_C}{2} \sum_p ||T_i \circ \vec{q}_i^{hp} - T_j \circ \vec{k}_j^{hp}||^2))$

    # Apply weighted attention.
    $o_i^h = \sum_j a_{ij}^h v_j^h$
    $\vec{o}_i^{hp} = T_i^{-1} \circ \sum_j a_{ij}^h (T_j \circ \vec{v}_j^{hp})$

    $s_i = \mathrm{Linear}(\mathrm{concat}_{h,p}(o_i^h, \vec{o}_i^{hp}, ||\vec{o}_i^{hp}||))$
    **return** $s_i$
**end procedure**

##### 2.4.2 Backbone Updating

The SwiftMHC Cross Structure module updates the backbone structure in two iterative blocks (**Algorithm 4, line 37**), taking the peptide representation $s_i$ as input. The backbone structure is initialized as identity frames, meaning the backbones of all peptide residues are overlapped at the origin and have the same orientations. Each block calls the *Backbone Update* module (**Algorithm 6**), that predicts an orientation quaternion $q_i$ and a translation vector $\vec{t}_i$. The two make up a frame $T_i$ that is to be transformed by the previous backbone frame, resulting in the next backbone frame: $T_i \leftarrow T_i \circ \text{BackboneUpdate}(s_i)$. This procedure is almost equal to **AlphaFold2**[1] **Suppl. Algorithm 23**, except that a quaternion is used for rotation, rather than a rotation matrix. The choice to use quaternions was made in order to reduce the number of floating point multiplications and additions to speed up the procedure.

**Algorithm 6** BackboneUpdate

**procedure** BackboneUpdate($s_i$):

  `# ` $b_i, c_i, d_i \in \mathbb{R}, \vec{t_i} \in \mathbb{R}^3$

  $b_i, c_i, d_i, \vec{t_i} = \text{Linear}(s_i)$

  `# Normalize the quaternion.`

  $q_i = (1, b_i, c_i, d_i)/\sqrt{1 + b_i^2 + c_i^2 + d_i^2}$

  $T_i = (q_i, \vec{t_i})$

  **return** $T_i$

**end procedure**

##### 2.4.3 Torsion Angles and Rigid Groups

SwiftMHC uses 8 frames to describe the conformation of every amino acid. The first frame is reserved for the orientation and the position of the backbone atoms: Cα and N, O and Cβ. The 7 remaining frames have been assigned to torsion angles and the corresponding rigid groups (**Fig. 2C** and **Suppl. Fig.7**). The selection of atoms for those rigid groups is the same as displayed in **AlphaFold2**[1] **Suppl. Table 2**.

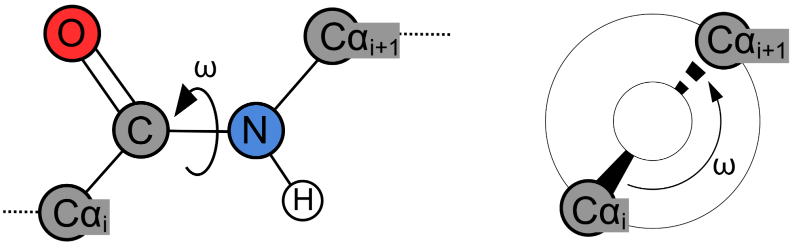

**Supplementary Figure 7: The ω torsion angle for the peptide bond.** A peptide bond between amino acid residue i and i+1 is shown (left). The ω torsion angle is measured from the positions of $C\alpha_i$ and $C\alpha_{i+1}$ around the C-N bond. A Newman projection of that same ω angle is also shown (right).

The ω torsion angle is a special case. In proteins, the bond between the carbonyl carbon (C) and the nitrogen (N) of the next amino acid is typically planar due to resonance, and the ω angle around this bond is often restricted to 0° (cis) or 180° (trans), with the trans configuration being much more common. After the backbone updating is complete and the positioning and orientation of atoms Cα, C and N in the backbone rigid groups is known, the ω angle is calculated using **Algorithm 7**. This ω angle is combined with the other predicted torsion angles (ϕ, ψ, $\chi_1$, $\chi_2$, $\chi_3$, $\chi_4$) to serve as input to the torsion loss function. This allows the network model to learn to avoid incorrect ω conformations.

**Algorithm 7** CalculateTorsion

**procedure** CalculateTorsion ($\vec{x}_1, \vec{x}_2, \vec{x}_3, \vec{x}_4$ ):

`# Calculate 3 interatomic bond vectors from 4 atomic positions.`

$\vec{b} = \vec{x}_3 - \vec{x}_2$

$\vec{n} = \frac{\vec{b}}{||\vec{b}||}$

$\vec{v}_1 = \vec{x}_2 - \vec{x}_1$

$\vec{v}_2 = \vec{x}_3 - \vec{x}_4$

`# Convert the bond vectors to Newman projections`

`# along a plane perpendicular to the central (2,3) bond.`

$\vec{p}_1 = \vec{v}_1 - (\vec{n}^\top \vec{v}_1)\vec{n}$

$\vec{e}_1 = \frac{\vec{p}_1}{||\vec{p}_1||}$

$\vec{p}_2 = \vec{v}_2 - (\vec{n}^\top \vec{v}_2)\vec{n}$

$\vec{e}_2 = \frac{\vec{p}_2}{||\vec{p}_2||}$

`# Calculate the torsion angle sin,cos from the Newman projections.`

$\alpha^{cos} = \vec{e}_1^\top \vec{e}_2$

**if** $(\vec{e}_1 \times \vec{e}_2)^\top \vec{n} < 0$ **then**

$\quad \alpha^{sin} = -||\vec{e}_1 \times \vec{e}_2||$

**else**

$\quad \alpha^{sin} = ||\vec{e}_1 \times \vec{e}_2||$

**end if**

$\alpha = (\alpha^{sin}, \alpha^{cos})$

**return** $\alpha$

**end procedure**

Newman projections represent the conformation of a molecule, by projecting the positions of the atoms around a specific bond in a plane, perpendicular to that bond (see **Suppl. Fig. 7** for an example). Such a projection provides insights into the torsion angle. The ω calculation function in **Algorithm 7** creates a Newman projection of the Cα atoms around each peptide bond: C-N. This is the type of bond that connects every pair of adjacent residues (**Suppl. Fig. 7**). From the resulting Newman projections, the *sin(ω)* and *cos(ω)* are calculated to be used in the loss function (**Subsection 2.6**).

We chose to calculate the cosines and sines of the torsion angles instead of the actual angles for two key reasons. First, this approach simplifies the torsion angle loss function, allowing the use of a straightforward normalization and squared error function when working with sin, cos vectors. Second, it improves computational efficiency, as cosines and sines can be directly computed using dot and cross products, eliminating the need for additional computationally expensive arcsin and arccos operations.

##### 2.5 Module 4: Residue-wise Binding Affinity Predictor

To enable SwiftMHC to handle peptides of varying lengths, we implemented a BA predictor that processes input sequences residue by residue (**Algorithm 8**). This BA predictor is called within the main module (**Algorithm 1, line 37**). Numerical BA is predicted by a multi-layer perceptron (MLP), that individually processes each residue of the updated sequence $\{s_i\}$. The BA prediction is the sum of the MLP outputs over all peptide residues (**Algorithm 8, line 8**). This BA value is trained to approach *1 - $log_{50000}(K_d)$* or *1 - $log_{50000}(IC_{50})$*, depending on which of the two data is available from the IEDB, where $K_d$ and $IC_{50}$ are experimentally determined parameters that reflect binding interactions and functional inhibition, respectively.

**Algorithm 8** Binding Affininty Predictor

1: **procedure** BindingAffinintyPredictor($\{s_i\}$):
2: # $i$ corresponds to a peptide residue.
3: # $r$ is the peptide length
4: # $i \in \{0, 1, 2, .., r\}$
5: $p_i = \text{Linear}(\text{relu}(\text{Linear}(s_i)))$
6:
7: # BA corresponds to $1 - \log_{50000}(\text{IC}_{50})$ or $1 - \log_{50000}(\text{K}_\text{d})$
8: $\text{BA} = \sum_i^r p_i$
9: **return** BA
10: **end procedure**

##### 2.6. Loss Terms

We train the modules in SwiftMHC from the following loss terms: *BA loss, backbone and side chain Frame Aligned Point Error (*FAPE*), Torsion angle loss* and *Structural Violations*. This subsection will explain each of these loss terms in detail.

*BA loss* ($\mathcal{L}_{BA}$ in **Algorithm 9**) is calculated on numerical output BA values that are compared in a MSE function against either the *1 - $log_{50000}(IC_{50})$* or *1 - $log_{50000}(K_d)$*. The BA loss value expresses how much the BA output deviates from the experimental value.

We also calculated FAPE loss as in AlphaFold2. FAPE measures the predictive error of the position of each atom $i$ relative to the local frame of residue $j$ (see **Suppl. Algorithm 28** and **Fig. 3f** in the AlphaFold2 paper for details). For example, if we hope to measure how well the CA of a peptide residue $i$ is predicted relative to a MHC residue $j$, we can calculate FAPE using: $d_{ij} = \sqrt{\epsilon + || T_j^{-1} \circ \vec{x}_i - T_j^{-1} \circ \vec{x}_i^{true} ||^2}$, where $\vec{x}_i$ is the predicted position for the CA atom of MHC residue $i$, $\vec{x}_i^{true}$ is the true position of the CA atom of peptide residue $i$, $T_j$ is the local frame of MHC residue $j$, $T_j^{-1}$ is the global to local frame transformation of residue $j$ and $\epsilon$ = $10^{-4}$ $Å^2$. The distance $d_{ij}$ is also clamped here using the same 10.0 Å distance. Using the local frames, FAPE also allows us to measure the chirality errors. Our FAPE loss contains a Backbone FAPE term and a side-chain FAPE term.

*Backbone FAPE* ($\mathcal{L}_{FAPE}^{backbone}$ in **Algorithm 9**) is calculated on predicted vs. true peptide Cα positions relative to the MHC structure. The MHC input frames $T_j$ and the true peptide positions $\vec{x}_i^{true}$ are derived from the target pMHC structure, while the input positions $\vec{x}_i$ are predicted from the peptide. This approach allows FAPE to measure the peptide's positions relative to the MHC it binds to. The resulting FAPE value expresses how different the output peptide frame positions are from the true peptide frames, as seen from inside each of the MHC frame-local coordinate systems.

*Side chain FAPE* ($\mathcal{L}_{FAPE}^{sidechain}$ in **Algorithm 9**) is calculated on the peptide side chain atoms, as described in **AlphaFold2**[1] **Suppl. Algorithm 28**. For this side chain loss term, both the input frames $T_i$ and input positions $\vec{x}_i$ are solely derived from the peptide, with no involvement from the MHC. This FAPE value expresses how different the atoms from the predicted peptide side chain are from their true positions, as seen from inside each of the peptide backbone-local coordinate systems. The true structural data originates from X-ray crystallography, where N, C, and O atoms are often indistinguishable. As a result, some amino acids exhibit 180° symmetry in their side chains, leading to the possibility of two valid frames. We use an implementation of the *renameSymmetricGroundTruthAtoms* algorithm (**AlphaFold2**[1] **Suppl. Algorithm 26**) to address this. Similar to AlphaFold2, a clamped L1-loss is applied to both backbone and sidechain FAPE (**AlphaFold2**[1] **Suppl. Algorithm 28, Line 4**).

*Torsion angle loss* ($\mathcal{L}_{torsion}$ in **Algorithm 9**) is calculated as described in **AlphaFold2**[1] **Suppl. Algorithm 27**. Only torsion angles (ϕ, ψ, ω, $\chi_1$, $\chi_2$, $\chi_3$, $\chi_4$) from the peptide are used in the equation, while MHC is ignored. The errors express how much these angles deviate from the angles in the true structure. For the side chain torsion angles in some amino acids there is a 180° symmetry, meaning that there can be two true angles. The torsion angle loss algorithm is designed to take both possibilities into consideration and select the best match.

**Algorithm 9** Calculate Loss

**procedure** CalculateLoss (BA, $\text{BA}^{true}$,
$\{\vec{x}_i^{all}\}, \{\vec{x}_j^{all}\}, \{\vec{x}_i^{true,all}\}, \{\vec{x}_i^{alt\ truth,all}\}$,
$\{T_i^{all}\}, \{T_j\}, \{T_i^{true,all}\}, \{T_i^{alt\ truth,all}\}$,
$\{\vec{\alpha}_i^{all}\}, \{\vec{\alpha}_i^{true,all}\}, \{\vec{\alpha}_i^{alt\ truth,all}\}$):
  `#` $i$ `corresponds to a peptide residue.`
  `#` $j$ `corresponds to a MHC residue.`

  `# For ComputeFAPE, see Alphafold2`[1] `Suppl. Algorithm 28`
  $\{\vec{x}_i^{N}, \vec{x}_i^{C\alpha}, \vec{x}_i^{C}, ..\} = \vec{x}_i^{all}$
  $\{\vec{x}_i^{true,N}, \vec{x}_i^{true,C\alpha}, \vec{x}_i^{true,C}, ..\} = \vec{x}_i^{true,all}$
  $\mathcal{L}_{\text{FAPE}}^{backbone} = \text{ComputeFAPE}(\{T_j\}, \{\vec{x}_i^{C\alpha}\}, \{T_j\}, \{\vec{x}_i^{true,C\alpha}\})$
  $\mathcal{L}_{FAPE}^{sidechain} = \text{ComputeFAPE}(\{T_i^{sidechain}\}, \{\vec{x}_i^{all}\}, \{T_i^{true,sidechain}\}, \{\vec{x}_i^{true,all}\})$

  `# For renameSymmetricGroundTruthAtoms, see`
  `# Alphafold2`[1] `Suppl. Algorithm 26`
  $\{T_i^{true,sidechain}\}, \{\vec{x}_i^{true,all}\} \leftarrow \text{renameSymmetricGroundTruthAtoms}($
  $\{T_i^{sidechain}\}, \{\vec{x}_i^{all}\}, \{T_i^{true,sidechain}\}$,
  $\{T_i^{alt\ truth,sidechain}\}, \{\vec{x}_i^{true,all}\}, \{\vec{x}_i^{alt\ truth,all}\})$

  `# For TorsionAngleLoss, see Alphafold2`[1] `Suppl. Algorithm 27`
  $\mathcal{L}_{torsion}^{all} = \text{TorsionAngleLoss}(\{\vec{\alpha}_i^{all}\}, \{\vec{\alpha}_i^{true,all}\}, \{\vec{\alpha}_i^{alt\ truth,all}\})$

  `# For StructuralViolations, see Algorithm 10`
  $\mathcal{L}_{\text{viol}} = \text{StructuralViolations}(\{x_i^{all}\}, \{x_j^{all}\})$

  `#` $\text{BA}^{\text{true}} \in \mathbb{R}$
  $\mathcal{L}_{\text{BA}} = \text{MeanSquareError}(\text{BA}, \text{BA}^{\text{true}})$

  `# Add on losses, according to tuning settings.`
  $\mathcal{L}_{\text{tot}} = 0$
  **if** baTune **then**
    $\mathcal{L}_{tot} += \mathcal{L}_{\text{BA}}$
  **end if**
  **if** fapeTune **then**
    $\mathcal{L}_{\text{tot}} += \mathcal{L}_{\text{FAPE}}^{backbone} + \mathcal{L}_{\text{FAPE}}^{sidechain}$
  **end if**
  **if** torsionTune **then**
    $\mathcal{L}_{\text{tot}} += \mathcal{L}_{\text{torsion}}^{all}$
  **end if**
  **if** fineTune **then**
    $\mathcal{L}_{\text{tot}} += \mathcal{L}_{\text{viol}}$
  **end if**
  **return** $\mathcal{L}_{\text{tot}}$
**end procedure**

*Structural violations* ($\mathcal{L}_{viol}$ in **Algorithm 9 & 10**) are calculated by comparing the predicted bond angles, bond lengths and non-bonded distances to literature values as described in **AlphaFold2**[1] **Suppl. Subsection 1.9.11**. For calculating bond angle and bond distance violations, only the peptide atoms are used. For clash violations, atoms from both the peptide and the MHC are used. The structural violation loss terms express how much the bond angles, bond lengths and non-bonded distances deviate from their literature values. Such loss terms are only included during the final *fine tuning* phase of training (**Subsection 3.1**).

**Algorithm 10** Structural Violations

**procedure** StructuralViolations ($\{x_i^{all}\}, \{x_j^{all}\}$):

    # Structural violations, computed in a similar way to
    # AlphaFold2[1] Suppl. Subsection 1.9.11

    # $i$ corresponds to a peptide residue.
    # $j$ corresponds to a MHC residue.

    # Bond length loss is calculated over backbone atoms only.
    # $l_{lit}^{general} = 1.329\text{Å}, l_{lit}^{proline} = 1.341\text{Å}$
    # $\tau^{bondlength} = 12\sigma, \sigma^{general} = 0.014\text{Å}, \sigma^{proline} = 0.016\text{Å}$
    $\mathcal{L}_{viol}^{bondlength} = \frac{1}{N_{bonds}} \sum_{i=1}^{N_{bonds}} max(\mid l_{pred}^i - l_{lit}^i \mid - \tau^{bondlength}, 0)$

    # Angle loss is calculated over backbone atoms only.
    # $\alpha_{lit}^{CNC\alpha} = 121.352°,\ \alpha_{lit}^{C\alpha CN} = 116.568°,$
    # $\tau^{bondangle} = 12\sigma, \sigma^{CNC\alpha} = 0.0311°, \sigma^{C\alpha CN} = 0.0353°$
    $\mathcal{L}_{viol}^{bondangle} = \frac{1}{N_{angles}} \sum_{i=1}^{N_{angles}} max(\mid cos\ \alpha_{pred}^i - cos\ \alpha_{lit}^i \mid - \tau^{bondangle}, 0)$

    # Clash loss is calculated between residues only
    # $\tau^{clash} = 1.5\text{Å}$
    # $\mathrm{N_{nbpairs}}$ counts the total number of non-bonded atom-atom interactions
    $\mathcal{L}_{\mathrm{viol}}^{clash} = \frac{1}{\mathrm{N_{nbpairs}}} \sum_{i=1}^{\mathrm{N_{nbpairs}}} \max(d_{lit}^i - d_{pred}^i - \tau^{clash}, 0)$

    $\mathcal{L}_{\mathrm{viol}} = \mathcal{L}_{\mathrm{viol}}^{bondlength} + \mathcal{L}_{\mathrm{viol}}^{bondangle} + \mathcal{L}_{\mathrm{viol}}^{clash}$
    **return** $\mathcal{L}_{\mathrm{viol}}$
**end procedure**

#### 3. Training Regimen

SwiftMHC was trained in PyTorch[3] 2.3.1 using the Adam optimizer, with a batch size of 16 and a learning rate of $10^{-3}$. To avoid large loss spikes, gradient norms are clipped to 0.5 at all times. Training is done in two phases (**Subsection 3.1**), where early stopping can end a phase when stopping criteria are met. In each phase, the early stopping conditions are reset and the model with the lowest validation loss is selected as the initial state in the next training phase.

##### 3.1 Training Phases

The training process consists of two distinct phases, each with specific settings (**Suppl. Table 1**). The goal of the first phase is to enable the model to learn structure prediction and reduce peptide Cα-RMSD from approximately 8 Å to 1-2 Å. During this phase, FAPE and torsion loss are included to guide atomic positions and torsion angles toward those of the true structure. Simultaneously, the BA loss term is incorporated to train the BA prediction module and to speed up the convergence of the 3D structure training. While this phase includes a high maximum epoch count (1000 epochs), an early stopping condition is applied to conclude the phase early if no more improvement in performance is observed (**Subsection 3.2**).

In the second phase, the model's structure prediction is fine tuned by introducing violation loss terms during backpropagation (**Subsection 2.6**). This phase resembles the fine tuning phase in AlphaFold2[1] and the associated loss terms encourage interatomic distances, bond lengths and bond angles to approach standard literature values.

**Supplementary Table 1: parameter settings for each training phase**
Columns 3-6 indicate which of the four loss terms are included in the back propagation during each phase (**Algorithm 9, lines 29-43**). Early stopping is turned on for both phases.

| phase | max epochs | FAPE loss | Torsion angle loss | violation losses | BA loss |
|---|---|---|---|---|---|
| 1 | 1000 | yes | yes | no | yes |
| 2 | 100 | yes | yes | yes | yes |

##### 3.2 Early Stopping

During training, an early stopping mechanism is applied to end a training phase and skip to the next phase or to end the training when the conditions are met in the final phase. The conditions for early stopping are based on loss values that are calculated on a validation dataset.

After each epoch the algorithm evaluates the loss values within a range of epochs (i.e., the patience, **Suppl. Fig. 8**) unless the number of epochs passed in that training phase is less than the patience value. We configured this patience value to be 50. For example, at epoch 50 we name the range 0 - 50 the *Frame of Patience*. At epoch 51, this frame shifts to the range 1 - 51 and so on.

Sometimes the high spikes of losses interrupts the early stopping. So we need to first smooth out the loss values within each Frame of Patience. Within each Frame of Patience the algorithm searches for outliers. Whether a loss value is an outlier or not is determined by the interquartile range (IQR). A loss value is considered an outlier if it lies more than 1.5 times the IQR above the third quartile (Q3) or more than 1.5 times the IQR below the first quartile (Q1). These outliers are replaced by the median value within the Frame of Patience, so that they cannot block early stopping.

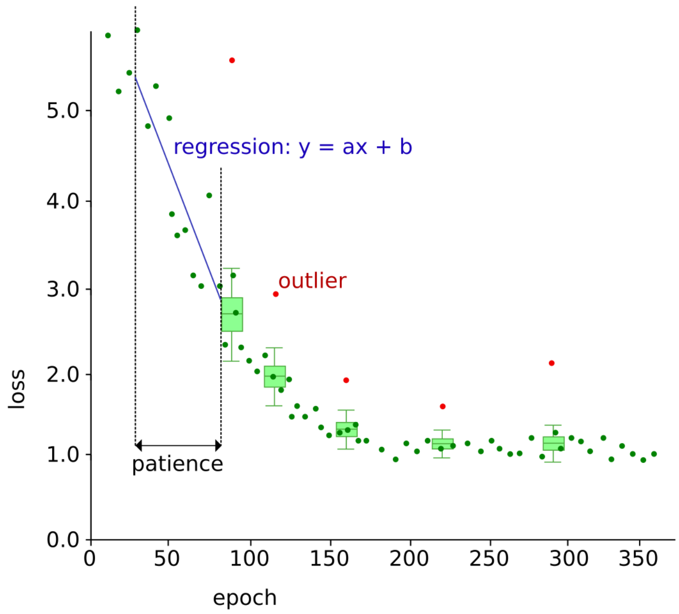

**Supplementary Figure 8: Early stopping criteria.** An example scatter loss plot is shown. Dots (dark green) indicate loss values. Box plots (light green) determine which dots are outliers (red) within the Frame of Patience (50 epochs). The best line (blue) is drawn between the non-outlier values by means of linear regression within the Frame of Patience. The parameters of this line are the slope (a) and offset (b).

After replacing the outliers with median values, linear regression is performed to find the best straight line through the loss values within a Frame of Patience. The algorithm stops a training phase early if the slope of this line is nearly horizontal and its value lies between $-1 \times 10^{-4}$ and $1 \times 10^{-4}$, indicating the predictive performance on the validation set is not improving any more.

#### 4. Supplementary Data

This section provides additional Ramachandran plots for backbone φ and ψ torsion angles of 3D models generated by SOTA approaches.

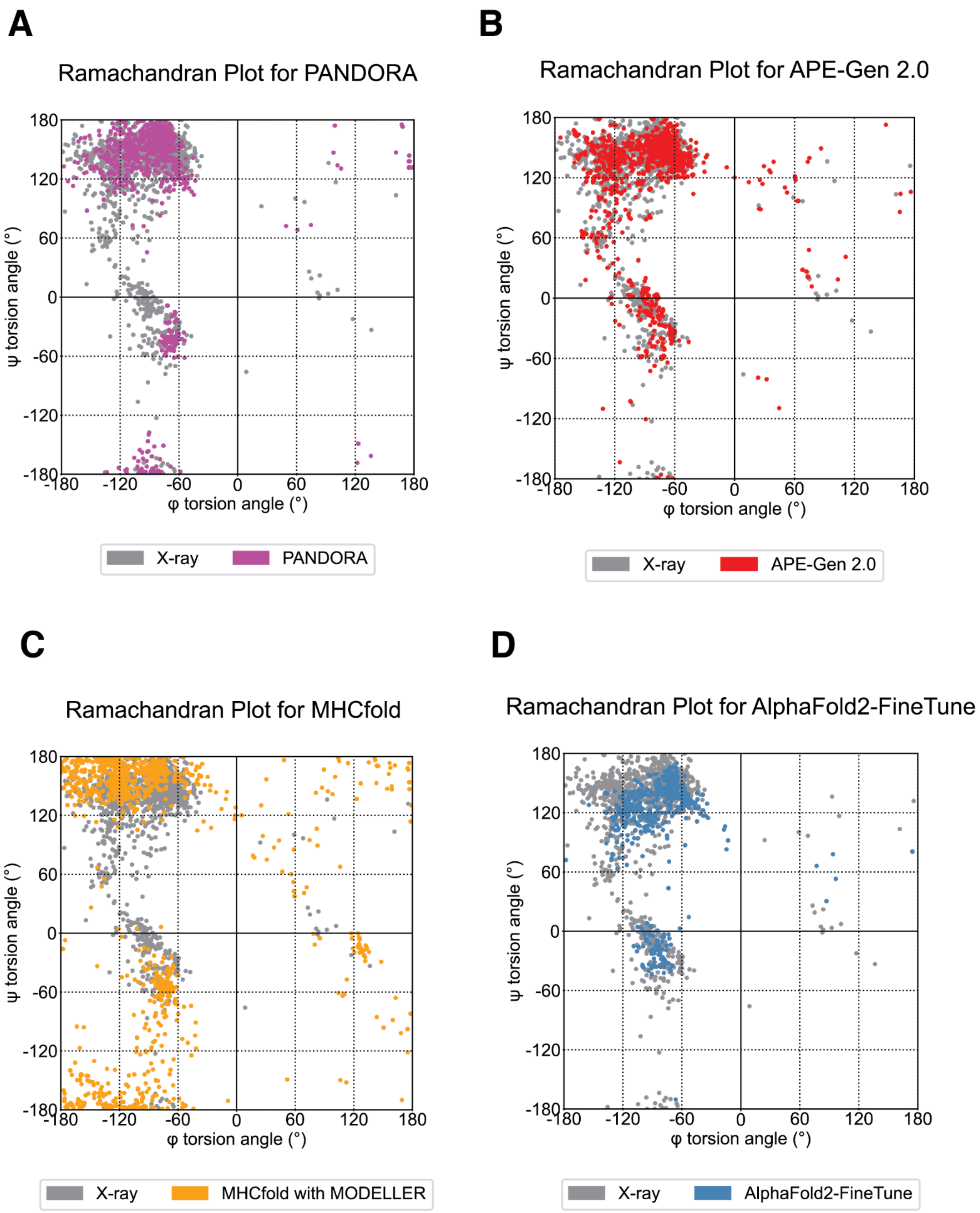

**Supplementary Figure 9. Distribution of backbone φ and ψ torsion angles of 3D models generated by SOTA approaches for the 202 X-ray test cases.** The plots have been overlaid with φ and ψ torsion angle data from the 202 X-ray structures. **A.** PANDORA models **B.** APE-Gen 2.0 models **C.** MHCfold models **D.** AlphaFold2-FineTune models

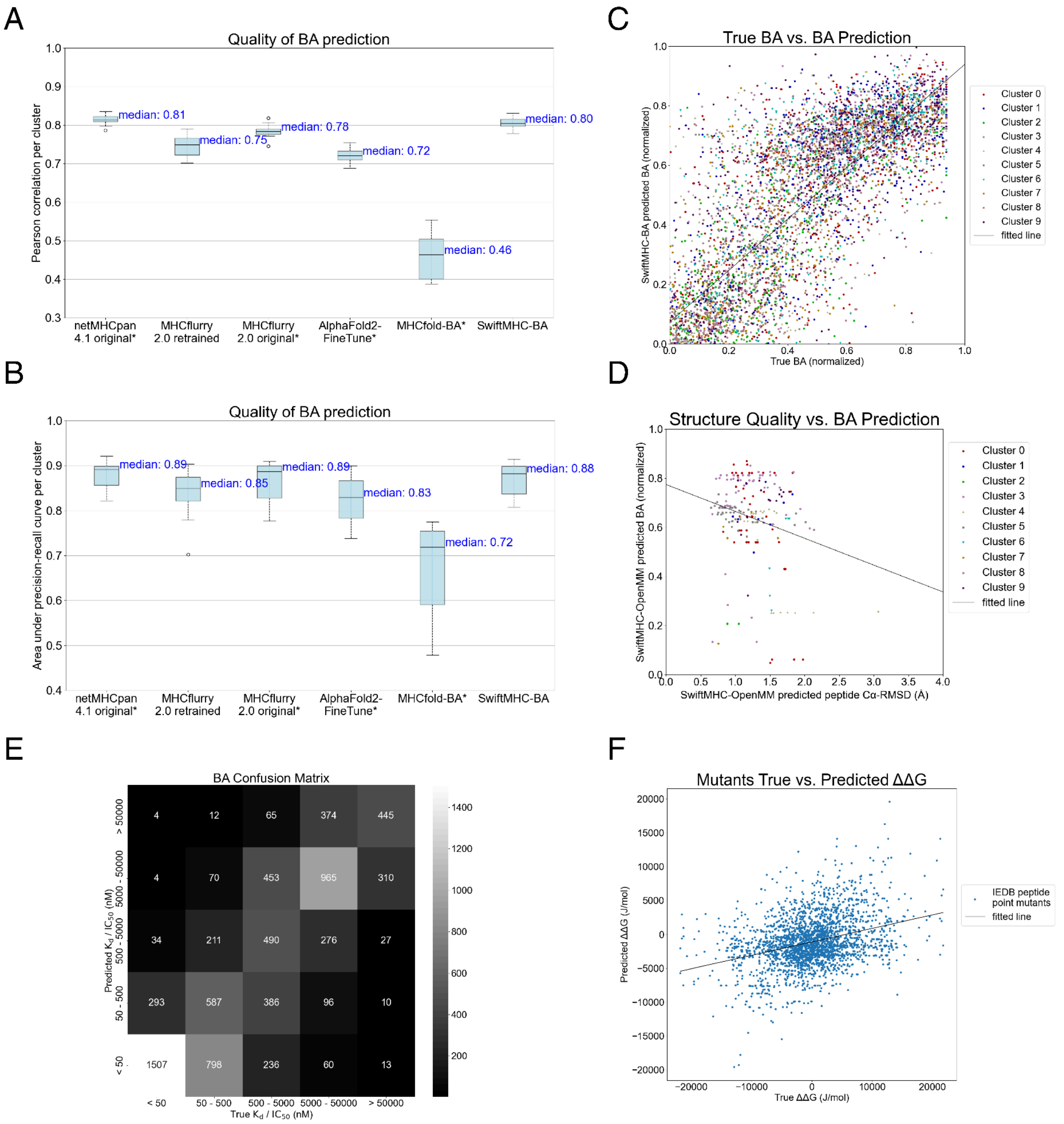

**Supplementary Figure 10. BA prediction metrics, correlation and training/test set overlap A.** BA prediction performance measured by Pearson correlation. Box plots show the distribution of Pearson correlation coefficients across 10 clusters of HLA-A02:01 9-mer peptides. **B.** BA prediction performance measured by area under the precision-recall curve (AUPR). Box plots show the distribution of AUPR across 10 clusters of HLA-A02:01 9-mer peptides. **C.** Scatter plot between SwiftMHC predicted BA and true BA for the 7,726 IEDB benchmark cases. The line of best fit was computed using least squares regression, with a pearson correlation of 0.82. BA values on the both the horizontal and vertical axes represent BA values, calculated as $1.0 - \log_{50000}(K_d)$ or $1.0 - \log_{50000}(IC_{50})$. **D.** Scatter plot between predicted BA and backbone RMSD for the 202 X-ray benchmark cases. The line of best fit was computed using least squares regression, with a pearson correlation of 0.22. BA values on the vertical axis represent SwiftMHC predictions, calculated as $1.0 - \log_{50000}(K_d)$ or $1.0 - \log_{50000}(IC_{50})$. **E.** Confusion matrix showing the 10-fold test set BA prediction distribution in the low and high affinity ranges. **F. Correlation between true and predicted ΔΔG for 2,838 single-point mutants of 9-mer peptides.** Each data point represents the change in **ΔG**, calculated as -RT $\log(K_d)$ or -RT $\log(IC_{50})$, with R=8.314 J $K^{-1}$ $mol^{-1}$ and T=298.15 K. The change in **ΔG** (Δ**ΔG**) is displayed per mutant, relative to its wild-type peptide. True $K_d$ or $IC_{50}$ data originates from the 7,726 data points taken from the IEDB. Predictions were made using SwiftMHC models trained on the same data fold as the corresponding wild-type. The Pearson correlation between true and predicted Δ**ΔG** values is 0.33.

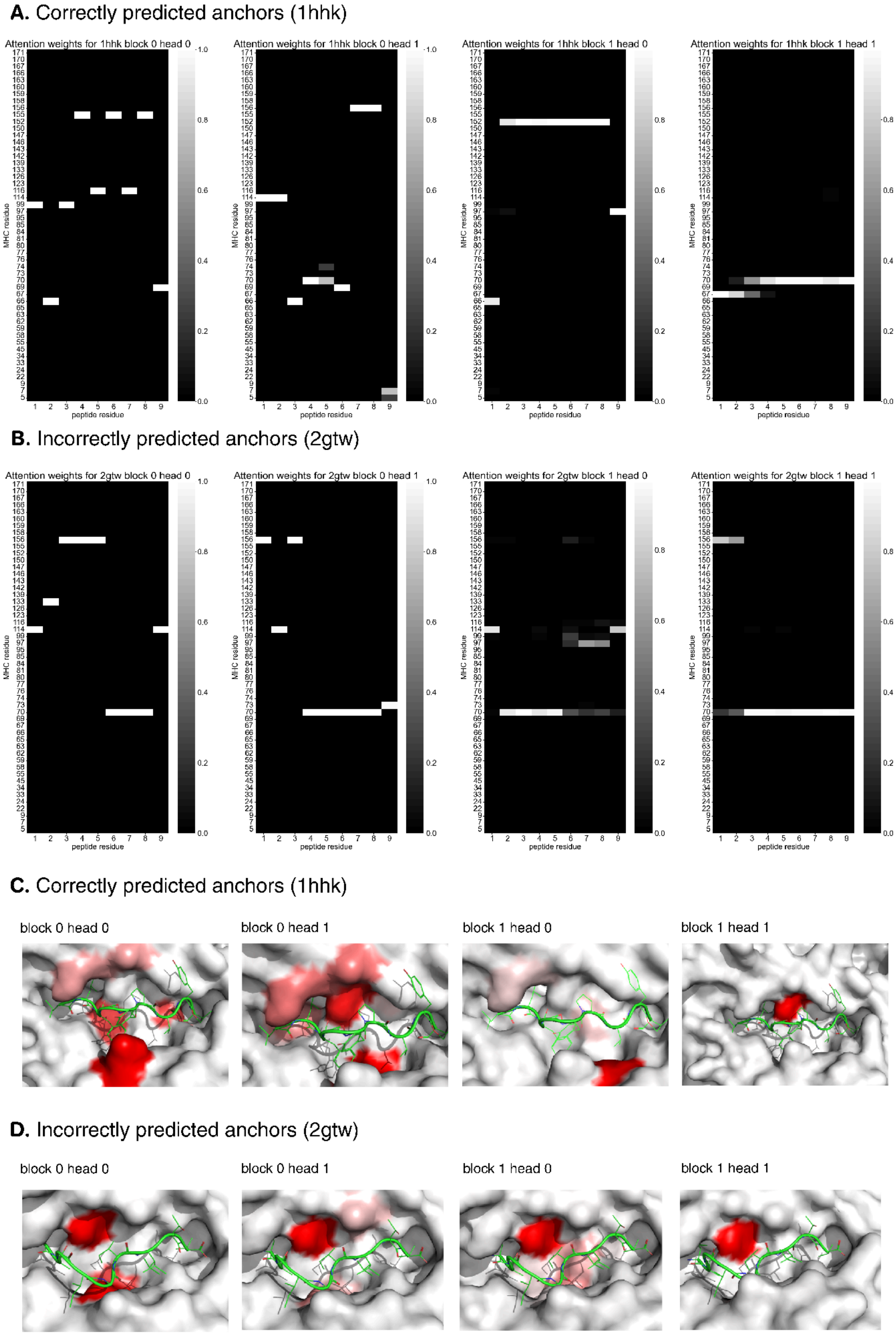

**Figure 11. Visualization of cross-attention weights in SwiftMHC. A.** Attention weight matrices from two transformer blocks (two heads each) for the 1HHK structure (peptide: LLFGYPVYV; allele: HLA-A02:01; X-ray anchor residues: positions 2 and 9), for which SwiftMHC correctly predicted the anchors **B.** Attention weight matrices for the 2GTW structure (peptide: LAGIGILTV; allele: HLA-A02:01; anchor residues: positions 1 and 9, for which SwiftMHC wrongly predicted the anchors **C and D.** Attention weights on the surface of the MHC structure of 1HHK and 2GTW, with regions colored in red according to the normalized sum of attention weights (each value was divided by the max sum of all MHC residues, darker red indicates higher attention). The peptide has been colored green. Notably, the

network correctly focused on the P2 pocket in 1HHK when the anchor residues are accurately predicted (**C**), whereas such focus on anchor pockets is absent in 2GTW, whose anchor was wrongly predicted (**D**). This indicates that attention patterns may provide a signal for whether the anchors are correctly predicted, although systematic analysis is needed to confirm this.

##### 4.1 SwiftMHC's robustness to structural variation in MHC molecules

We considered how best to evaluate SwiftMHC's robustness to structural variation in MHC molecules. Although the reviewer suggested assessing homology modeling variance, we reasoned that X-ray structures are already used as templates in such models and therefore directly analyzed X-ray structures instead. Specifically, we performed an all-against-all TM-align comparison of the G-domains (IMGT residues 2–180) from 202 HLA-A*02:01 X-ray structures. The pairwise RMSD values showed a mean of **0.53 Å** and a maximum of **1.19 Å**, indicating minimal structural variability within this allele. Given this limited range, and considering that the training models were themselves constructed using the homology models from X-ray templates, we did not pursue additional robustness experiments. We therefore believe SwiftMHC should be reasonably robust to homology-modeled MHC structures.

##### 4.2 Edge cases on SwiftMHC's vs. sequence-based approaches

When evaluating binary classification performance (binder vs. non-binder using a 500 nM threshold), we found 315 cases where the retrained MHCflurry 2.0 correctly predicted binding status and SwiftMHC did not, and 508 cases where SwiftMHC was correct and MHCflurry 2.0 was not. See **Fig. 12AB** for the sequence logo representations of the corresponding peptides. No substantial differences were observed between these two sets of peptides.

When evaluating numerical binding affinity prediction (normalized, as 1.0 - $\log_{50000}(K_d)$ or 1.0 - $\log_{50000}(IC_{50})$ ), we found that in 2977 cases the retrained MHCflurry 2.0 was closer to the true value and in 4749 cases SwiftMHC was closer. The mean absolute distance to the normalized true value was 0.18 for MHCflurry and 0.14 for SwiftMHC. See **Fig. 12CD** for the sequence logo representations of the corresponding peptides. No substantial differences were observed between these two sets of peptides.

Therefore, we are not able to identify a pattern when the sequence-based approaches outperform SwiftMHC.

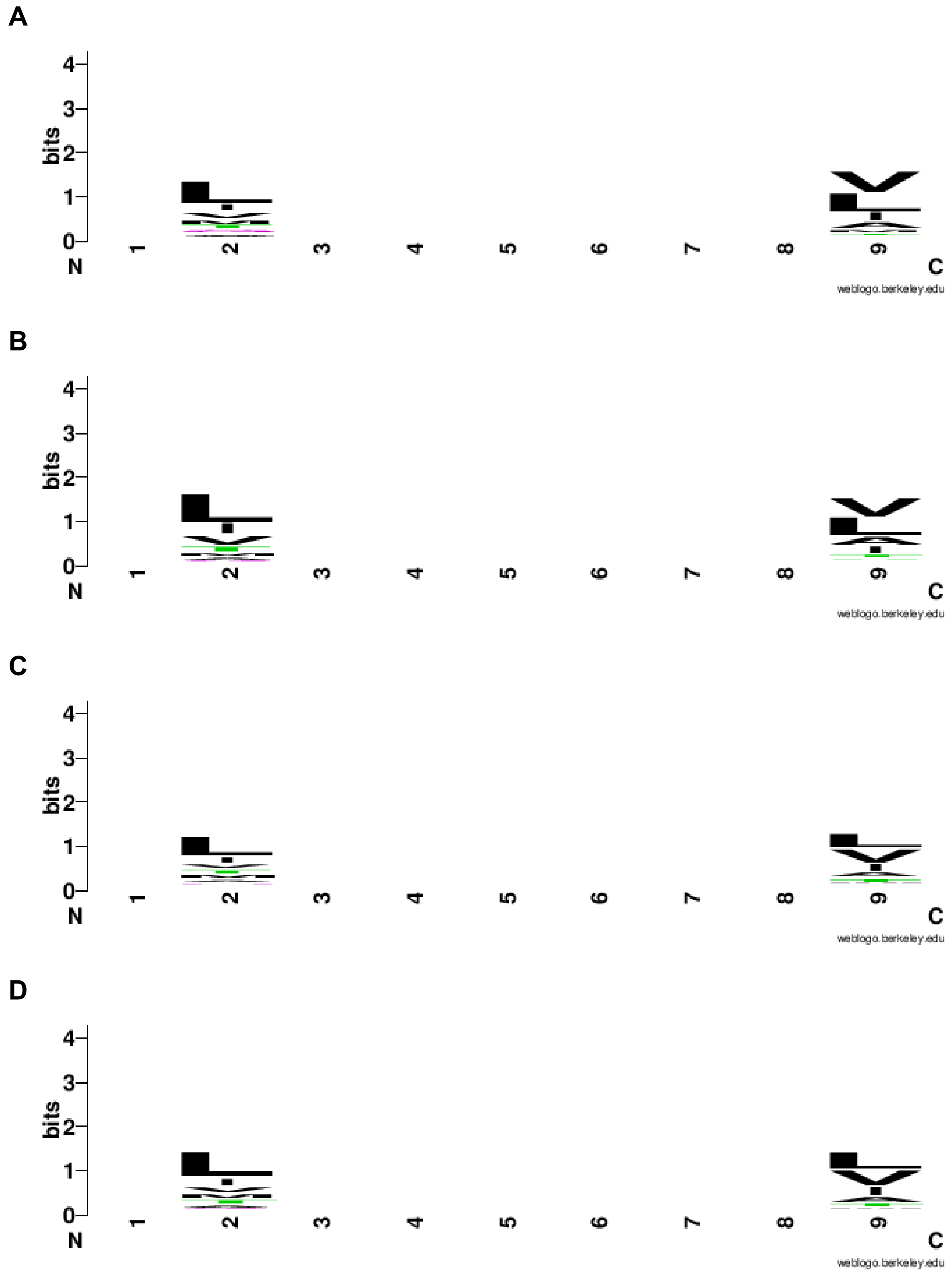

**Figure 12**. **Sequence logos for sets of peptides that were scored by SwiftMHC and retrained MHCflurry 2.0.** These images were generated by weblogo (*https://weblogo.berkeley.edu/logo.cgi*). **A.** representation for 508 cases where SwiftMHC was right about classifying a peptide as binding/non binding and MHCflurry was wrong. **B.** representation for 315 cases where MHCflurry 2.0 was right about classifying a peptide as binding/non binding and SwiftMHC was wrong. **C.** representation for 4749 cases where SwiftMHC's BA prediction was closer to the true BA value than MHCflurry 2.0's. **D.** representation for 2977 cases where MHCflurry's BA prediction was closer to the true BA value than SwiftMHC's.
